# Hierarchical cross-linking of a bacterial spore coat Hub protein

**DOI:** 10.64898/2026.02.03.703564

**Authors:** Khira Amara, Catarina Fernandes, David Q.P. Reis, Daniela Silva, Carmen Olivença, Bruno Gonçalves, Tiago Pais, Maria L. Martins, Guillem Hernandez, Cristina Timóteo, Ricardo A. Gomes, Ana S. Pina, Mónica Serrano, Tiago N. Cordeiro, Adriano O. Henriques

## Abstract

Hub proteins are highly connected nodes in protein-protein interaction networks and are often intrinsically disordered proteins (IDPs) or contain intrinsically disordered regions. In *Bacillus subtilis*, the morphogenesis of the spore surface is orchestrated by a set of so-called morphogenetic proteins that guide the assembly of distinct layers. Formation of the inner coat is directed by SafA^FL^ and its shorter isoform, C30. Both are expressed early in sporulation under the control of σ^E^ and localize at the interface between the developing inner coat and the underlying cortex peptidoglycan. From this site, they act as organizational hubs, recruiting client proteins essential for coat maturation. Among these is Tgl, a transglutaminase synthesized later in development following activation of σ^K^ after engulfment completion. We show that the C30 domain exhibits IDP-like features yet self-assembles into >1200 kDa complexes stabilized by disulfide bonds and that these bonds are required for subsequent proper Tgl-mediated “spotwelding” cross-linking. Small-angle X-ray scattering (SAXS) and photobleaching show that Tgl immobilizes but does not drastically alter these assemblies. These findings support a hierarchical, biphasic model for inner coat assembly: initial self-assembly and disulfide stabilization, followed by Tgl-mediated cross-linking and structural stabilization. According to this model, the forms of SafA^FL^/C30 that dominate the two stages recruit different client proteins in register with the course of morphogenesis.

## INTRODUCTION

Highly connected Hub proteins are central nodes in interactomes and play essential roles in the organization and connectivity of many cellular components (1–3). The Hub proteins are often intrinsically disordered proteins (IDPs) or have disordered regions (IDRs) which allows them to bind multiple partners (1–3). Static Hubs bind their partners simultaneously at different docking sites while in dynamic Hubs a single site may competitively bind to short linear motifs (SLiMs) often located in IDRs of several client proteins; these may include protein-modifying enzymes, that compete for the same site in the Hub (1–3). Proteins, so-called morphogenetic, that intervene in the morphogenesis of the bacterial spore, arguably the most resistant cellular entity known to us, govern the assembly of different structural sub-layers and, as we claim, behave like Hubs.

In the model organism *Bacillus subtilis* spores are encased into several protein layers, differentiated into a lamellar inner layer, a more external, striated and electrodense outer layer, in turn surrounded by a glycosylated crust (4–6). Spore differentiation takes place in a sporangium that consists of a larger mother cell and a smaller, polar forespore, or future spore (**Fig 1A**). Formation of the coat and crust layers of spores relies on the synthesis of over 80 proteins in the mother cell, under the control of transcription factors σ^E^, first, and α^K^, later (**Fig 1A**) (5,6). These proteins are targeted to their final sub-layers, as morphogenesis progresses, by a group of proteins known as morphogenetic (4–6). During the early stages of coat and crust assembly, the morphogenetic proteins localize to the surface of the developing spore, to form organizational centers that recruit the components of the different layers (7). SafA governs inner coat assembly, while CotE guides assembly of the outer coat, and CotZ is the main organizer of crust assembly (8) (**Fig 1B**). Recruitment of these proteins to the spore surface requires two other proteins, SpoVM and SpoIVA (**Fig 1B**). SpoVM is a 26 amino acids founder protein that recognizes the positive curvature of the forespore whereas SpoIVA is an ATPase that covers the spore surface with cables formed as the result of ATP-dependent polymerization forming a basal scaffold for assembly of the rest of the coat (9–11). Following the initial localization step, the coat and crust proteins eventually cover the entire surface of the spore in a process called encasement that calls for the function of another morphogenetic protein, SpoVID (12) (**Fig 1B**).

**Fig 1.**
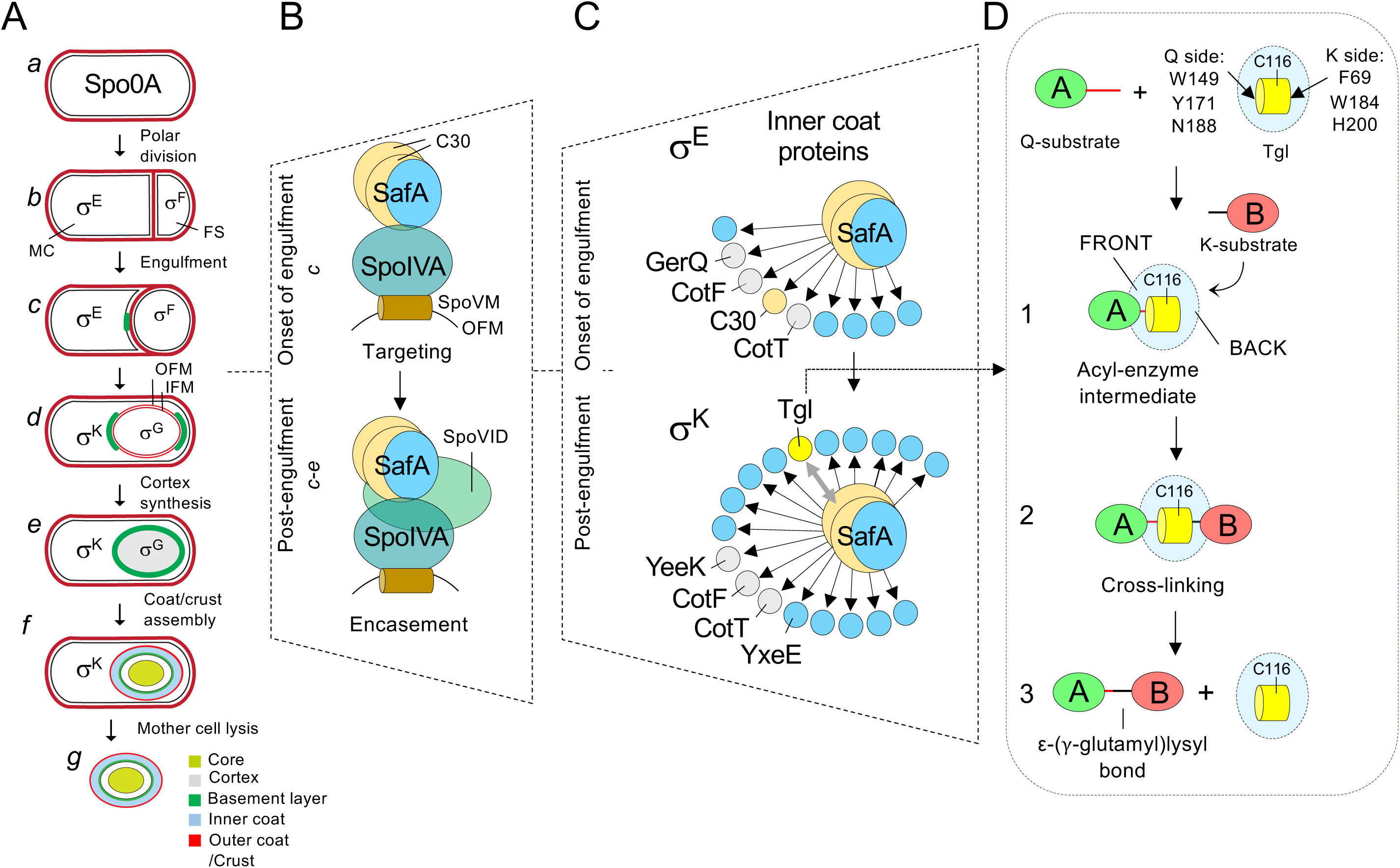
Role of SafA in spore coat assembly. **A:** Main stages of sporulation: pre-divisonal cells (*a*); sporangia that have undergone polar division (*b*), at the beginning of engulfment (*c*), following engulfment completion (*d*), during the last stages in synthesis of the spore cortex and the coat and crust layers (*d*-*f*) and when spores are released (*g*). The localization of SafA-YFP is represented in green (panels *c* to *e*). The cell type-specific sigma factors and their window of activity is indicated. **B**: In a targeting step, soon after engulfment begins, SafA is recruited to the forespore by the SpoIVA ATPase. SpoIVA localization at or close to the forespore outer membrane in turn relies on SpoVM. In a second step, encasement of the forespore by SafA depends on an interaction with SpoVID. **C**: SafA is the Hub for inner coat assembly; it recruits proteins produced both under the control of σ^E^ and σ^K^, such as Tgl. Assembly of SafA is auto-regulatory as the recruitment of Tgl results in cross-linking of SafA (two way arrow). CotE, required for outer coat assembly and CotZ, required for crust assembly, are not represented for simplicity. **D**: Tgl reaction cycle. In a first step, a Gln donor substrate approaches the enzyme from the Q-side and forms an acyl-enzyme covalent intermediate. In a second step, a Lys substrate (acyl acceptor) engages the enzyme from the K-side and the cross-link is formed. The reaction occurs in a hydrophobic tunnel that harbors the catalytic residues including Cys116. The ceiling of the tunnel is thought to open to release the cross-linked product. The mostly hydrophobic residues that line the Q- and K-side entries of the tunnel are indicated.

Morphogenetic proteins SafA, CotE and CotZ function as Hubs that recruit the inner and outer coat/crust proteins. SpoVID, in turn, is a Hub that coordinates encasement by the inner coat outer coat/crust modules (13). This study deals with the assembly pathway of SafA, the Hub for inner coat assembly. The *safA* mRNA is produced early in the mother cell, under σ^E^ control (14) but because of alternative translation initiation at an internal methionine codon, it yields three proteins: the full-length (SafA^FL^, 43 kDa; migrates at about 45 kDa in SDS-PAGE), C30 (25 kDa; migrates at 30 kDa), corresponding to the C-terminal domain of SafA^FL^, and SafA^N21^ (18 kDa, migrates at 21 kDa) which corresponds to the N-terminal moiety of SafA^FL^ (15) (**Fig 2A**). SafA^FL^ has a LysM domain in its N-terminal end which is both a protein-protein and a protein-peptidoglycan interaction domain (**Fig 2A**) (16). Early in development, the LysM domain together with a region termed A, form a localization signal required for an interaction first with SpoIVA, which targets SafA^FL^ to the spore surface, then with SpoVID, which allows encasement (12,17–20). At a late stage in development, the LysM domain is involved in the interaction of SafA^FL^ with the spore cortex peptidoglycan, which is formed underneath the coat (16). Immunogold labeling showed that SafA localizes at the cortex/inner coat interface suggesting a role as a molecular staple linking the two structures (16,17). Spores of a *safA* deletion mutant lack the morphological features associated with the inner coat and lack several proteins (17,21). These are the SafA client proteins shown, using GFP fusions, to localize to the inner coat, consistent with the Hub model (8,22). Importantly, it is C30, and the C30 domain of SafA^FL^ that are responsible for the recruitment of the inner coat proteins (23). Since C30 has no localization signal, its targeting to the spore surface requires the interaction with the C30 domain of SafA^FL^ (24). The C30 domain also interacts strongly with Itself (24).

**Fig 2.**
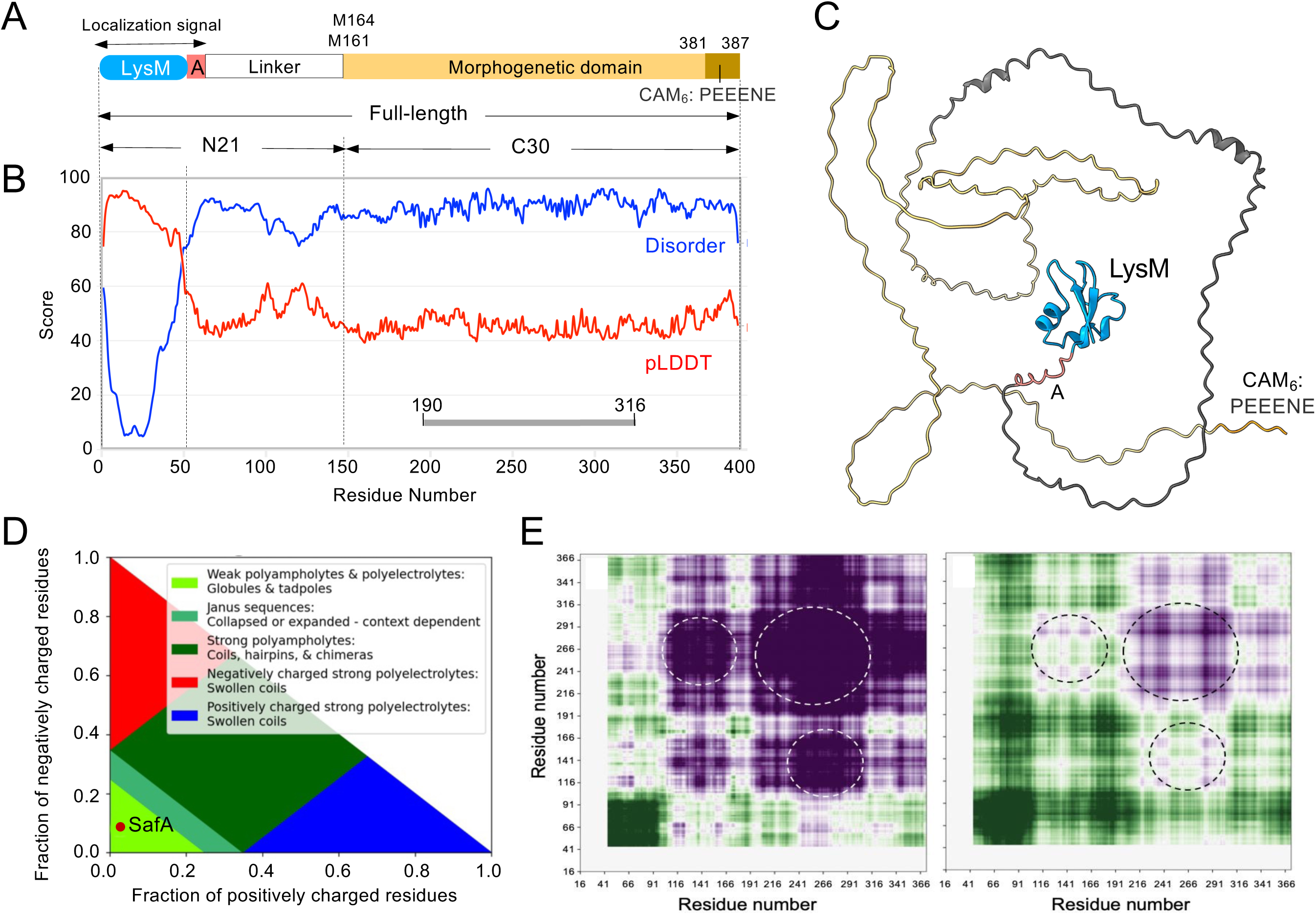
Domain organization and properties of SafA. **A**: SafA^FL^ has an N-terminal LysM domain which together with a region termed A forms a localization signal. This signal is required for targeting SafA to the spore surface via an interaction with SpoIVA, and for encasement, through an interaction with SpoVID. The LysM domain is also important for the interaction of SafA with the cortex peptidoglycan while C30 is the morphogenetic domain of the protein, responsible for the recruitment of the inner coat proteins. C30 is also made independently through internal translation of the *safA* mRNA initiating at methionine codons 161 or 164. A third form of the protein, SafA^N21^, which includes the LysM domain, region A and the linker, has no known function. A 6-mer short acidic peptide (CAM_6_) at the C-terminal of SafA^FL^ or C30 is also indicated. C30 relies on an interaction with the C30 domain of SafA^FL^ for its localization to the coat. **B**: SafA is an intrinsically disordered protein (IDP). Disorder propensities, calculated using Metapredict (blue) (51) are compared to the structural confidence calculated by AlphaFold 2 (AF-2; red), as measured by the predicted local distance difference test (pLDDT). The two predictions are in anti-phase: Metapredict indicates high disorder in regions where AlphaFold assigns low structural confidence. These findings support a model in which SafA has a well-structured LysM domain, followed by highly disordered regions, particularly in region A and the C30 domain. **C**: Structure of SafA predicted with AF-3. The left panel shows the structure colored based on SafA’s molecular architecture (LysM, region A, linker, C30, and CAM_6_, as in panel **A**). **D**: Diagram of states partitioned into five distinct conformational classes based on the fractions of positively and negatively charged residues. SafA is found within the region associated with compact conformations, suggesting that it may adopt a more condensed, globule-like structure despite its disordered nature. **E:** Self-intermolecular interaction maps (inter-maps) between the intrinsically disordered regions (IDRs) of SafA reveal potential sub-regions of intermolecular interactions. These interactions were predicted using the chemical physics encoded within the Mpipi-CG (52) and CALVADOS 2.0 (53) force fields using FINCHES (Ginell et al., 2025). Attraction and repulsion are displayed using a purple-to-green gradient, indicating varying interaction strengths within the IDRs, particularly in the C30 region. Both force fields predict attractive interactions within residues the stretch defined by residues 191-316 (this region is represented as a gray line in panel **B**).

One of the proteins that SafA^FL^/C30 recruits to the forespore is a transglutaminase, Tgl, which catalyzes the formation of ε-(γ-glutamyl) lysyl cross-links between protein-bound glutamyl and lysil residues (23,25) (**Fig 1C** and **D**). Tgl controls the assembly and/or extractability of at least four proteins involved in spore coat assembly. These proteins are more extractable from spores of a *tgl* deletion or a catalytic mutant and are thought to be direct substrates of Tgl. SafA^FL^, C30 and GerQ are produced early, under σ^E^ control while YeeK is also produced late, under σ^K^ control (**Fig 1C**) (21,22). They all localize to the inner layers of the coat (8,22). Tgl is produced under the control of α^K^ and it also localizes to the inner layers of the coat in a manner that requires interactions with its pre-assembled substrates (23,25,26) (**Fig 1C**). Tgl has a hydrophobic tunnel that shields the catalytic triad from water; one of the entrances of the tunnel is the glutamyl (Q) acceptor site, the other is the lysil acceptor site (25,27). In the reaction cycle, an acyl-enzyme intermediate is formed with the Q substrate; the enzyme then engages the K substrate and the cross-link is produced (**Fig 1D**).

Remarkably, the single alanine substitutions that impair the most the localization of Tgl are those of residues that contour the Q acceptor entry of the tunnel, followed by substitutions on the K side, and finally substitutions of the active site residues (25) (**Fig 1D**). Thus, Tgl is directed to its coat-localized substrates in a way that promotes formation of the acyl-enzyme intermediate, the first step in the reaction. Assembly of SafA^FL^/C30 is auto-regulatory in that recruitment of Tgl leads to the cross-linking of the two proteins; at least SafA^FL^ is cross-linked by Tgl in both the cortex and coat layers of the spore (23) (**Fig 1C**). C30 forms oblong hexamers in solution, as determined by small angle X-ray scattering (SAXS) (25) but how these oligomers are formed and how they are cross-linked by Tgl is unknown. Tgl, however, is postulated to exert a spot welding cross-linking activity fortifying the pre-assembled complexes of C30 and its other substrates at the cortex/inner coat interface (25).

As we now show, C30 is cross-linked by Tgl and is the only domain of SafA^FL^ cross-linked by the enzyme *in vitro*. As such, we examined the assembly pathway of C30, as a proxy for the assembly of SafA^FL^. We show that the previously observed C30 hexamer (C30_6_) is most likely the unit for the self-assembly of higher order complexes. We show that disulfide bonds contribute critically to the proper assembly and stability of SafA-containing complexes and are essential for efficient C30 cross-linking by Tgl later in development. This defines a hierarchical cascade of protein-protein cross-linking events. SAXS analysis reveals that Tgl-mediated cross-linking does not induce significant changes in the overall shape or size of the complexes, consistent with the spot-welding cross-linking of pre-assembled complexes. The SafA assembly pathway is biphasic: during early development, under the control of σ^E^, self-association and disulfide bond formation predominate; later, following engulfment completion and activation of σ^K^, Tgl is expressed, recruited to the spore surface by its pre-assembled substrates, and catalyzes their cross-linking in situ to reinforce the cortex–inner coat interface.

## RESULTS

### The structural organization of SafA

SafA^FL^ exhibits a modular architecture; it consists of an N-terminal LysM domain that together with a short stretch of 12 residues termed region A, forms a localization signal; a linker connects the localization signal to the C30 domain (**Fig 2A**, top; **S1A_Fig**). C30 ends with a 6-mer acidic motif, PEEENE, termed CAM_6_ (**Fig 2A**). C30, alone or in the context of SafA^FL^, is the morphogenetic domain responsible for the recruitment of the inner coat proteins (25). Structure predictions using AlphaFold (AF) (28,29) revealed a highly confidence structure for the LysM domain (pLDDT > 90) but failed to predict a defined structure for region A, the linker and C30, the last two of which show a high propensity for disorder (**Fig 2B** and **2C**). As such, SafA^FL^ classifies as an intrinsically disordered protein (IDP). Strikingly, however, when SafA^FL^ is subjected to limited proteolysis with Trypsin or AspN, a fragment corresponding to at least part of the C30 domain accumulates, as confirmed by N-terminal analysis, despite the presence of two potential canonical trypsin cleavage sites and several AspN sites within the domain (**S1_Fig**). This indicates a resistance to proteolysis that is atypical for IDPs. BSA, a globular protein, also shows resistance to trypsin cleavage, whereas α-sinuclein, a well characterized IDP is eventually degraded by trypsin (**S1B_Fig**). Part of the explanation for the proteolysis resistance comes from sequence analysis, which shows that the disordered regions downstream of the LysM domain can be classified as “globule” (**Fig 2D**) (30). These “globule” regions adopt more compact yet flexible conformations that might confer resistance to proteolysis (**Fig 2D**). Moreover, within the C30 region, several potential attractive self-interaction hotspots are predicted (31,32) (**Fig 2E**, regions encircled; see also **S1_Table**). The presence of these hotspots is consistent with the strong self-interactions of SafA^FL^ (via the C30 domain), of C30 alone, and of SafA^FL^ with C30, as detected in a yeast two hybrid assay (17) and with the formation of a C30 hexamer in solution as detected by size exclusion chromatography (SEC) and small-angle X-ray scattering (SAXS) (25). We presume that oligomerization of SafA^FL^ through the C30 domain contributes to protease resistance. Interestingly, Ozin and co-authors noticed the similarity of C30 to human amelogenin, the predominant matrix protein involved in enamel formation and also an IDP (17,33–35). The similarity encompasses a Pro-rich stretch, residues M1 to S119 of C30 (161 to 280 in SafA^FL^) and residues I50 to D157 in amelogenin (**S2_Fig**). These residues in amelogenin are located in a central hydrophobic region (or HR) that contributes to the intermolecular interfaces formed during self-assembly of amelogenin into supramolecular structures, in turn important for enamel templating (33–35) (**S2_Fig**; see also the Discussion).

We have shown before that the activity of Tgl decreases with increasing concentrations of C30 because the enzyme becomes part of the cross-linked products and that this effect is suppressed by increasing enzyme concentration (25). We now show that these properties also apply to SafA^FL^ (**S3_Fig** and **S4_Fig**; see also **S1_Text**). Moreover, we show that in spite of the presence of several Lys and Gln residues in the LysM domain, region A and in the linker region, purified SafA^N21^, which encompasses these three regions (**Fig 2A** and **S1A_Fig)**, is not cross-linked by Tgl (**S5A_Fig**). Here, we used C30 as a proxy to study the Tgl-catalyzed cross-linking of the inner coat Hub.

### Disulfide bonds are required for the accumulation of C30 oligomers past a dimer

In the presence of the reducing agent dithiothreitol DTT, C30 forms an oblong hexamer in solution, C30_6_ (25). C30, however, has four cysteine (Cys) residues, which raises the possibility that disulfide bonds promote the formation and/or stabilization of C30_6_ or higher order oligomers (**Fig 3A**). C30 also harbors two lysine (Lys) residues, expected to be involved in the formation of ε-(γ-glutamyl)-lysyl cross-links catalyzed by Tgl; the role of these residues, however, has not been directly addressed (**Fig 3A**). The Cys residues in C30 are arranged into two groups. Each group consists of closely-spaced residues: one group, formed by Cys214 and Cys228, is located close to the N-terminal end of C30, and close to one of the two lysine residues, Lys177; the other group of Cys residues, formed by Cys323 and Cys325, is located towards the C-terminal end of the protein, and just downstream of the second Lys residue, Lys318 (**Fig 3A**; numbering refers to the position in SafA^FL^). To study the role of the Lys and Cys residues in the oligomerization and cross-linking of C30, we overproduced and purified several variants of C30 along with the WT protein, hereinafter C30^WT^: in one variant, the four Cys residues were substituted by serine (Ser) residues (dubbed C30^C to S^); in another variant, the two Lys residues, Lys177 and Lys318, were replaced by alanine (Ala) (C30^K to A^); a third variant had both the Lys and the Cys residues replaced (C30^K to A/C to S^). As assessed by reducing SDS-PAGE, C30^WT^ and C30^C to S^ run at about 30 kDa while the variants lacking the Lys residues (C30^K to A^ and C30^K to A/C to S^) migrate slightly slower (**Fig 3B**, left panel). Under non-reducing conditions, however, C30^WT^ runs as a main species close to the interface of the stacking and resolving gels and also as larger forms that remain at the well (**Fig 3B**, right panel). For reasons that will become apparent further down, the species at the stacking/resolving gel interface is most likely an oligomer formed by repeats of the C30_6_ hexamer, and is herein named (C30_6_)_n_; the form that stays at the well may consist of higher-order repeats of (C30_6_)_n_ and is designated as [(C30_6_)_n_]_n_.

**Fig 3.**
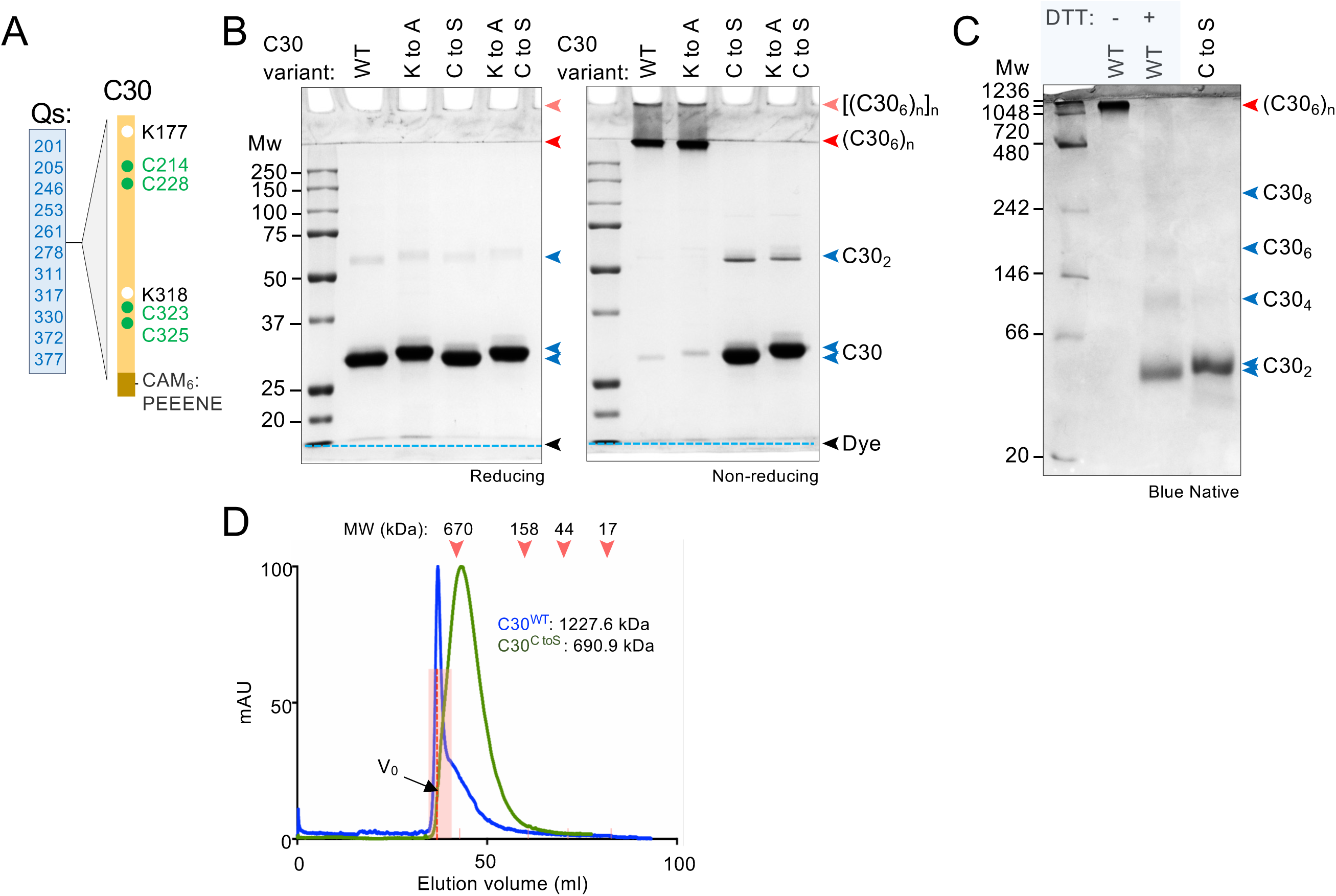
C30 forms disulfide cross-linked oligomers. **A**: Diagram of C30 showing the position of Cys (green dots) and Lys (white dots) residues (numbering is from the N-terminus of SafA^FL^; see also **S1_Fig**). **B**: C30^WT^, C30^K to A^ (the two Lys residues replaced by Ala), C30^C to S^ (the four Cys residues replaced by Ser) and C30^K to A/C to S^ (the two groups of substitutions combined) were resolved by SDS-PAGE (12.5% gels) under reducing (left panel) or non-reducing (right panel) conditions. The blue dashed lines shows the dye front. **C**: C30^WT^ and C30^C to S^ were resolved by BN-PAGE (5-15% gradient) in the absence or in the presence of DTT (-/+ 2 mM). In panels **B** and **C**, the position of the relevant proteins is indicated by arrowheads. The gels were stained with Coomassie. **D**: SEC analysis of C30^WT^ and C30^C to S^ on a Sephacryl S-300 column. The estimated sizes of C30^WT^ and C30^C to S^ are shown. The elution positions of blue dextran (void volume, V_0_) and of molecular weight standards (in kDa) are indicated.

Evidently, the Lys residues are not required for the oligomerization of C30^WT^ because (C30_6_)_n_ (at the resolving/stacking gel interface) and [(C30_6_)_n_]_n_ (at the well) are still detected for the C30^K to A^ variant in non-reducing SDS-PAGE (**Fig 3B**, right panel). In sharp contrast, neither (C30_6_)_n_ nor [(C30_6_)_n_]_n_ are detected for C30^C to S^ or C30^K to A/C toS^ in non-reducing SDS-PAGE (**Fig 3B**, right panel). Together, these observations suggest that C30 oligomers are stabilized by disulfide bonds. A possible dimer, C30_2_, is detected by reducing and non-reducing SDS-PAGE for both C30^C to S^ and C30^K to A/C to S^ (**Fig 3B**). Hence, a dimer may be an intermediate in the formation of (C30_6_)_n_. In C30^C to S^*, i.e*., in the absence of disulfide bonds, C30_2_ may also dissociate in the presence of SDS (**Fig 3B**; see also below).

As judged by Blue Native PAGE (BN-PAGE), in which oligomeric proteins are resolved in their native state primarily according to size (36,37), the C30^WT^ oligomer(s) are detected as a single species above the 1048 kDa marker band (**Fig 3C**). When C30^WT^ is pre-incubated with DTT, however, the complex dissociates into species with sizes close to those of C30_2_ (the most abundant species), C30_4_, C30_6_ and at least C30_8_ (the latter three species barely detected) (**Fig 3C**). Importantly, the profile obtained for the DTT-treated C30^WT^ is nearly identical to that obtained for C30 ^C to S^ (**Fig 3C**). Thus, in line with the results described above, and as judged by SDS- and BN-PAGE, this suggests that: *i*) disulfide bonds are only needed for the stabilization of C30 oligomers past C30_2_ at least under the conditions of BN-PAGE; *ii*) the formation of C30_6_ and higher order oligomers proceeds by the association of C30_2_ dimers (see more below related to this specific point); *iii*) the C30_2_ dimer, in turn, appears to be the smallest building block of C30_6_ and large oligomeric forms.

As a further test of these predictions, we analyzed C30^WT^ and C30^C to S^, which were purified in the absence of DTT, by size exclusion chromatography (SEC) using a Sephacryl S-300 column that was also run under non reducing conditions. C30^WT^ eluted mainly in a single peak, peak 1, close to the void volume of the column (**Fig 3D**). Since this peak elutes close to the upper resolution limit of the Sephacryl S-300 column (1500 kDa for globular proteins) its mass cannot be determined with high accuracy; with an estimated mass of 1227 kDa, it likely corresponds to a species containing between 6 and 8 hexameric units (expected mass 1080–1440 kDa; **Fig 3D**). This size range is consistent with the BN-PAGE analysis (**Fig 3C**; 1048 kDa) and suggests the presence of about 45 molecules of C30 in the oligomer. Nevertheless, the asymmetry of the SEC peak, skewed to the right, suggests that the main species is in an equilibrium with smaller forms (**Fig 3D**). In contrast, C30^C to S^ eluted as a main species of about 670 KDa, possibly containing 22 C30 molecules, (**Fig 3D**). The peak was broader as compared to C30^WT^ and also skewed to the right (**Fig 3D**). Possibly, the C30^C to S^ 670 kDa complex is also in an equilibrium with smaller species but is more promptly dissociated (**Fig 3D**). The comparison of the BN-PAGE, in which C30^C to S^ is detected mostly as a dimer, with the SEC data shows that large oligomers of C30^C to S^ exist in solution but do not withstand the conditions of BN-PAGE. Absent from C30^C to S^ complexes, disulfide bonds could allow C30 complexes to endure the conditions of electrophoresis. DDM can preserve stable complexes but not detergent-sensitive assemblies (36,37), suggesting that the self-assembly of C30 proceeds at least in part through hydrophobic interactions.

### Hydrophobic interactions contribute to the assembly of C30 oligomers

To directly test whether disulfide bonds allow complexes of C30^WT^, but not those of C30^C to S^, to endure the presence of DDM during BN-PAGE (**Fig 3C** and **3D**), we applied C30^WT^ and C30^C to S^, at a concentration of 0.3 mg/ml, to a Sephacryl S-300 column equilibrated with a buffer containing 0.05% DDM, the same concentrations used for BN-PAGE. C30^WT^ eluted as a main peak close to the 670 kDa marker (**S6A_ Fig**, peak 2, estimated mass of 810 kDa), with a trailing to the right, and a smaller peak (peak 3 at around 39 kDa). Presumably, not all of the molecules in the C30^WT^ complexes are disulfide cross-linked, and DDM can dissociate the uncross-linked molecules from the complexes. In addition, a subtle inflexion in the region of the chromatogram to the left of peak 2, overlapping with the void volume, was detected (**S6A_Fig;** peak 1, at around 1129 kDa). As expected, DDM partially dissociated C30^C to S^ which showed a peak close to the void volume of the column (**S6B_Fig**, peak 1, about 1175 kDa), a second peak at around 669 kDa (peak 2) and a third at around 43 kDa (peak 3). The 43 kDa peak most likely corresponds to the 39 species detected in the C30^WT^ chromatogram. The larger species (peaks 1 and 2) are not detected by BN-PAGE, in line with the idea that the electrophoresis conditions are more disruptive than those of SEC. In any event, the assembly pathway of C30 seems to involve hydrophobic interactions that are disrupted by DDM, and disulfide bonds that stabilize the complexes.

### C30 dimers have a parallel orientation within higher order oligomers

The arrangement of the four Cys residues (**Fig 3A** and **S1A_Fig**) raised the question of whether they were equally important for the assembly or stability of C30 oligomers. We generated C30 variants in which either the N-terminal Cys Cys214 and Cys228 or the C-terminal Cys323 and C325, were replaced by Ser. These forms of C30 are hereinafter designated C30^C to S N-ter^ and C30^C to S C-ter^. We first showed that in reducing SDS-PAGE, all the proteins migrate as monomers with traces of dimers (**Fig 4A**) and we then proceeded to dissect the role of the N- and C-terminally located Cys residues. C30^C to S N-ter^ was mainly detected as a dimer under non-reducing SDS-PAGE (although traces of a possible tetramer are seen) and as a tetramer in BN-PAGE (**Fig 4B** and **4C**). Thus, the absence of the N-terminal Cys residues impairs the ability of C30 to accumulate as complexes past C30_4_. Moreover, the tetramer is most likely a dimer of intramolecularly cross-linked dimers in which the disulfide bonds involve the C-terminal Cys residues (**Fig 4E**). We note, however, that although the main species detected by non-reducing SDS-PAGE appear to be dimers, monomers are also seen, which suggests that disulfide bonds may also form between each dimer pair (**Fig 4E**, designated intermolecular cross-links).

**Fig 4.**
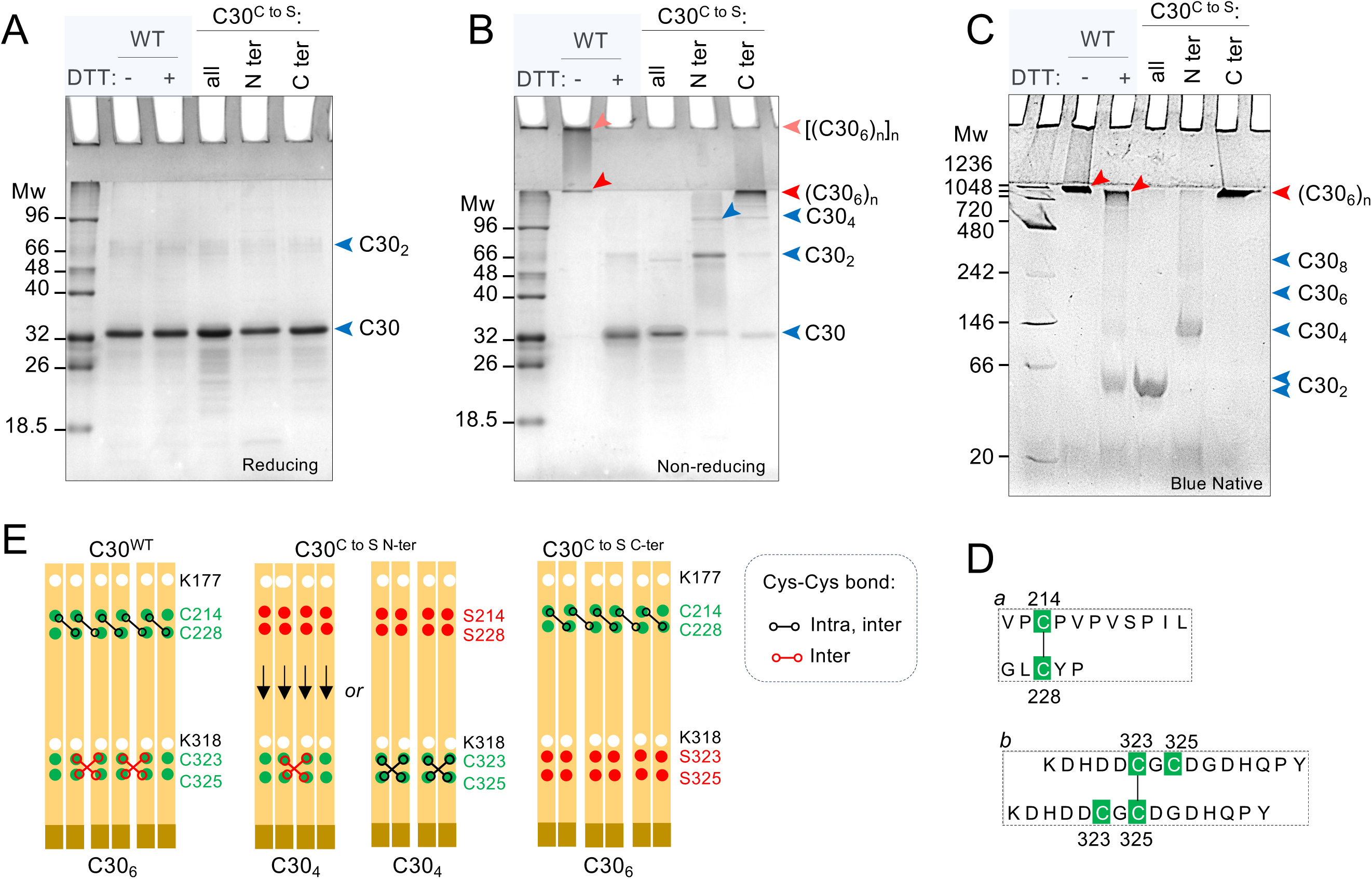
The N- and C-terminal pairs of Cys residues have different functions. **A** and **B:** C30^WT^, C30^N-ter^ (Cys214 and Cys228 replaced by Ser) and C30^C-ter^ (Cys323 and Cys325 replaced by Ser) and C30^C to S^ (all Cys residues replaced by Ser) were resolved by SDS-PAGE (12.5% gels) under reducing (**A**) or non-reducing (**B**) conditions. C30^WT^ was loaded in the absence (WT-DTT) and after treatment with DTT (2 mM; WT +DTT) **C**: The same proteins as in **A** were resolved by BN-PAGE (5-15% gradient) and the gels stained with Coomassie. C30^WT^ was also analyzed after incubation with DTT (as above). In **A** and **B**, the various oligomeric forms of C30 are indicated by arrowheads. The gels were stained with Coomassie. **D**: C30 has a parallel orientation within dimers (C30_2_) and in higher order complexes (C30_6_). Disulfide bonds involving the C-terminal Cys residues (red crosses) orient and stabilize the dimer, while the N-terminal Cys residues cross-link adjacent dimers (black). The sizes of the various species detected by different electrophoretic techniques is also show at the bottom of the panel. **E**: Mass spectrometry analysis of the C30_2_ dimer identifies disulfide bonds between Cys214 and Cys228 (*a*). and Cys323-Cys325 (*b*; see also **S7_Fig**).

On the other hand, C30^C to S C-ter^ is detected mainly as a band at the stacking/resolving gel interface, (C30_6_)_n_, both in non-reducing SDS-PAGE and in BN-PAGE (**Fig 4B** and **4C**). Therefore, Cys323-Cys325 are important for the accumulation of oligomers larger than (C30_6_)_n_ and in their absence the assembly pathway stops at (C30_6_)_n_ (**Fig 4B**). In contrast, the N-terminal Cys214 and Cys228 residues form disulfide bonds that are important for the accumulation of oligomers up to (C30_6_)_n_ and in their absence the assembly pathway stops at C30_4_ (above). Importantly, the detection of C30^C to S N-ter^ and C30^C to S C-ter^ cross-linked oligomers indicates that disulfide bonds form within (but not between) the N- and C-terminal groups of Cys residues and this in turn indicates a parallel orientation of the C30 monomers (**Fig 4D**). Consistent with this interpretation, mass spectrometry analysis revealed a Cys214-Cys228 cross-link in the (C30_6_)_n_ species at the stacking/resolving gel interface while a Cys323-Cys325 cross-link was detected for C30^WT^ (**Fig 4E** and **S7_Fig**; see also **S1_Text**). No cross-links were detected between Cys residues in the N-terminal group and those in the C-terminal group (**Fig 4E** and **S7_Fig**).

### Disulfide bonds allow for proper cross-linking of C30 by Tgl

Having shown the importance of the Cys residues for the self-assembly and stability of C30 oligomers, we wondered whether the disulfide bonds would also be important for their cross-linking by Tgl. First, C30^WT^ and C30^C to S^ were incubated for 120 min in the absence or in the presence of Tgl. C30^WT^ migrated at about 30 kDa in reducing SDS-PAGE and as a large species, (C30_6_)_n_, under non-reducing conditions, while C30^C to S^ is detected mainly at around 30 kDa in either case (**Fig 5A**). This is consistent with the requirement for disulfide bonds for the formation/stabilization of C30 oligomers (above). Following incubation with Tgl, both C30^WT^ and C30^C to S^ are detected close to the interface of the stacking/resolving gels, (C30_6_)_n_, and at the well, [(C30_6_)_n_]_n_ (but formation of this species from C30^C to S^ is inefficient) in both reducing or non-reducing conditions (**Fig 5A**). Thus, Tgl cross-links both C30^WT^ and C30^C to S^. The Tgl-cross-linked C30^WT^ complex shows the same apparent migration as the non-cross-linked complex detected under non-reducing SDS-PAGE (**Fig 5A**). This is in keeping with the idea that Tgl cross-links pre-assembled complexes (23,25). In the presence of Tgl, and given sufficient time, no C30 monomers remain detectable in the SDS-PAGE gel implying that in the larger oligomers, all of the C30 units are cross-linked. In contrast, the monomer of C30^C to S^ persists until 2 hours (**Fig 5A**). C30^C to S^ is cross-linked by Tgl, but the cross-linked products run slightly faster in SDS-PAGE than those of C30^WT^ under both reducing and non-reducing SDS-PAGE (**Fig 5A**, notice their position relative to the red line marking the stacking/resolving gel interface in the bottom panel). The behavior of C30^C to S^ alone or in the presence of Tgl does not seem to result from an indirect effect of the four Cys to Ser substitutions because C30^WT^ produced similar results when incubated with DTT prior to reaction with Tgl (**Fig 5A**). Strikingly, the cross-linking of C30^C to S^ by Tgl generated at least one species that migrates ahead of the 30 kDa position on the gel (**Fig 5A**, brown arrowhead). Because proteolysis seems unlikely, this species may represent a form of C30^C to S^ with an intramolecular cross-link. If so, with only one free Lys residue, the reaction with Tgl would only produce a cross-linked dimer (see also the following section). One role of the disulfide bonds in C30 may thus be to reduce the occurrence of unwanted cross-linking reactions.

**Fig 5.**
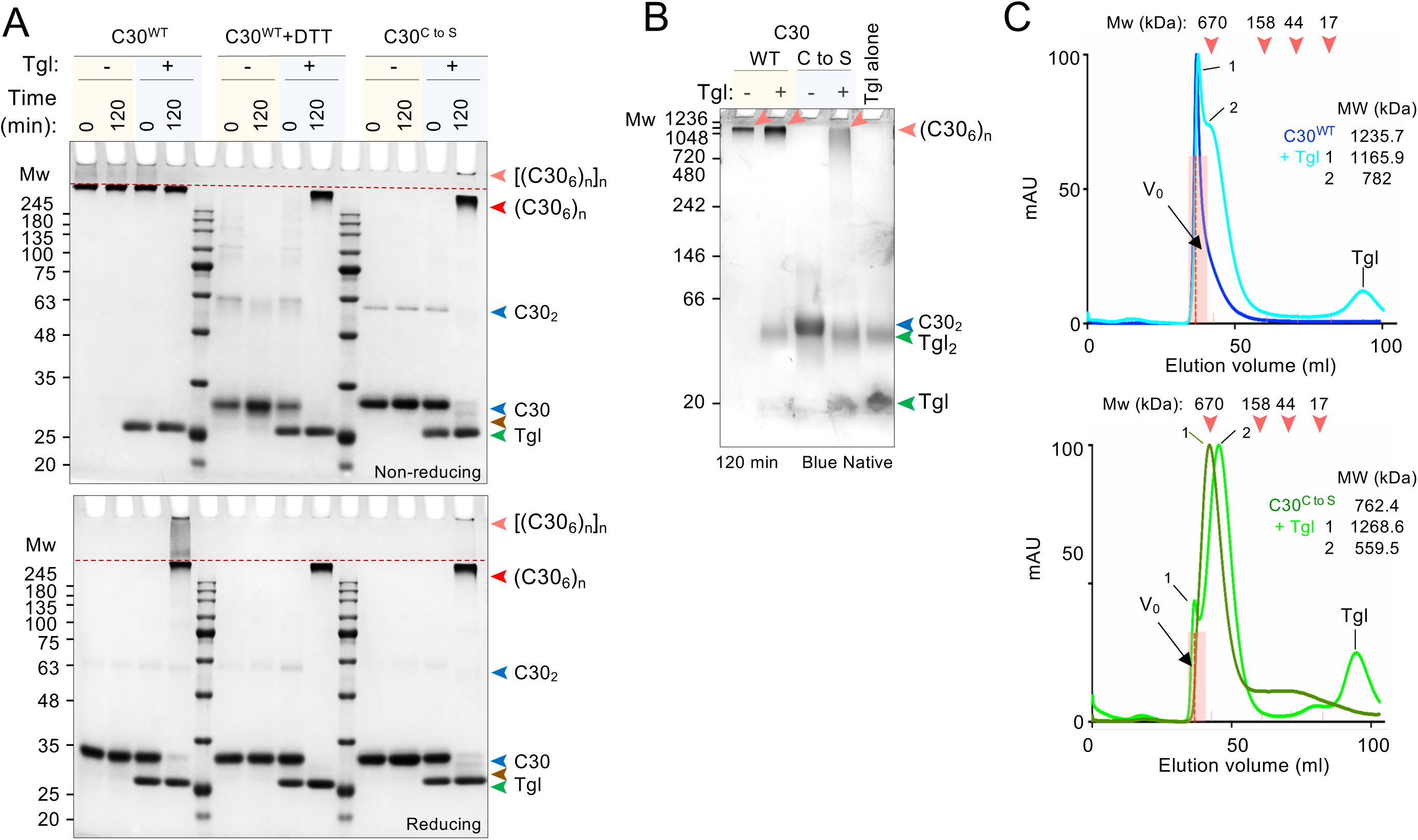
Disulfide bonds are required for the formation of large C30 oligomers. **A**: C30^WT^, pre-incubated in the absence or in the presence of DTT (2 mM) or C30^C to S^ were incubated for 120 min in the absence (“-“) or in the presence (“+”) of Tgl. The reaction products were resolved by non-reducing (upper panel) or reducing (lower panel) SDS-PAGE (10%) and the gels stained with Coomassie. **B**: C30^WT^ or C30^C to S^ were incubated alone (“-“) or with Tgl (“+”) for 120 min and then the proteins were resolved by BN-PAGE (5-15% gradient). In **A** or **B**, the position of C30 and C30_2_ are indicated by a blue arrowhead and the position of higher order oligomers is indicated by the red arrowheads. The position of Tgl is indicated by a green arrowhead and the brown arrowhead indicates a faster-migrating form of C30. **C**: SEC analysis of C30^WT^ (top panel) and C30^C to S^ (bottom), before and after incubation with Tgl, using a Sephacryl S-300 column. The estimated sizes of C30^WT^ and C30^C to S^ before and after cross-linking are shown. The elution positions of blue dextran (void volume, V_0_) and of molecular weight standards (in kDa) are indicated.

In BN-PAGE the cross-linked C30^C to S^ runs as a smear close to the top of the resolving gel in contrast with the sharp band of the cross-linked C30^WT^ (**Fig 5B**). This suggests that Tgl cross-links complexes of C30^C to S^ that are, on average, smaller than those formed by C30^WT^ (**Fig 5B**) as also noticed in the SDS-PAGE gels (**Fig 5A**), in line with the observation that the 762.4 kDa complex of C30^C to S^ detected by SEC may be in an equilibrium with smaller species (**Fig 3D**, above). The BN-PAGE also revealed a monomer and a dimer of Tgl (**Fig 5B**). A crystallographic Tgl dimer was also detected although the resolved structure was of a monomer (27).

The SEC analysis of the reaction products of Tgl with C30^WT^ shows a main peak of about 1165.9 kDa (**Fig 5C**, upper panel, peak 1) and a second peak of about 782 kDa (peak 2). Although peak 1 elutes close to the upper resolution limit of the Sephacryl S-300 column (1500 kDa for globular proteins) and its mass cannot be accurately estimated, it may represent a species with between 6 to 8 copies of an hexamer (expected mass 1080-1440 kDa). Peak 2 may correspond to a species with 4 copies of an hexamer (estimated 720 kDa). For C30^C to S^, a peak of 1268.6 kDa is detected, possibly with 6 to 7 copies of an hexamer (expected mass of 1092-1260 kDa) (**Fig 5C**, lower panel, peak 1). A more prevalent peak, corresponding to a mass of 559.5 kDa, could be a trimer of hexamers, whose expected mass is of 540 kDa (peak 2). We infer that the formation of disulfide bonds allows for the proper cross-linking of C30 oligomers by Tgl. Importantly, the complete conversion of C30^WT^ into large DTT and SDS-PAGE-resistant oligomers implies that all of the monomers or other building blocks in a complex must be cross-linked via the Lys (and Gln) residues. Whether cross-linking of C30 within the complexes follows a defined order is not yet known (but see **S1_Text** and **S8_Fig**).

### Disulfide bonds are important for the of C30 cross-linking by Tgl

The parallel arrangement of C30, the formation of Cys214-Cys228 and Cys323-Cys325 bonds and the proximity of Lys177 and Lys318 close to the N- or C-terminal groups of Cys residues suggested that the two types of cross-linking could be to some extent interdependent (**Fig 3A** and **S1_Fig**). The disulfide bonds could place Lys177 and Lys318 at the right distance and orientation for their efficient cross-linking to a Gln in an adjacent C30 molecule (**Fig 3A** and **S1_Fig**; eleven Gln residues are distributed along the entire length of C30). If so, then the kinetics of C30^C to S^ cross-linking could be impaired relative to the WT. We tested this idea by measuring the kinetics of cross-linking of different forms of C30. In control reactions in which C30^WT^ or C30^C to S^ were incubated without Tgl for up to 120 min, no high molecular weight species were detected (**Fig 6A** and **6B**; first three lanes of the panels). The substrate proteins were then incubated with Tgl (“+” signals) and samples of each reaction were collected over time and analyzed by reducing SDS-PAGE to follow the appearance of the cross-linked species. The activity of Tgl was monitored by quantifying the disappearance of C30^WT^ or C30^C to S^ over time. Three features distinguish the cross-linking of C30^WT^ and C30^C to S^. Firstly, the kinetics of cross-linking started to slow for C30^C to S^ after 15 min compared to C30^WT^, and unlike the latter, a fraction of non-cross-linked C30^C to S^ remains at the end of the experiment (**Fig 6**, compare panels **A** and **B**). Note that enzyme activity was quantified as the rate of disappearance of C30 relative to time 0 (**Fig 6E**) or the rate of formation of C30^WT^ or C30^C to S^ oligomers, of the (C30_6_)_n_ type, relative to time 0 (**Fig 6F**). Secondly, after 30 min the [(C30_6_)_n_]_n_ species starts to accumulate at the well for C30^WT^ but is barely detectable for C30^C to S^ (**Fig 6A** and **6B**). We infer that disulfide bonds are important for the formation of [(C30_6_)_n_]_n_. We note, however, that even for C30^WT^, (C30_6_)_n_ is not fully converted into [(C30_6_)_n_]_n_ during the duration of the experiment (**Fig 6A**), consistent with the SDS-PAGE (non-reducing conditions) and SEC analysis of C30^WT^ before and after reaction with Tgl (above; see **Fig 5A** and **5C**). Lastly, the form of C30^C to S^ with a possible intramolecular cross-link was also detected (**Fig 6B**, brown arrowhead; see also **Fig 5A** and text above). Importantly, Tgl interacts with C30^WT^ and C30^C to S^ with the same affinity (K_d_ around 0.05-0.07 µM), as assessed by microscale thermophoresis (**Fig 6G**). Thus, the differences in the cross linking of C30^WT^ and C30^C to S^ are likely to be directly related to catalysis and not to the recognition of the two substrates by the enzyme.

**Fig 6.**
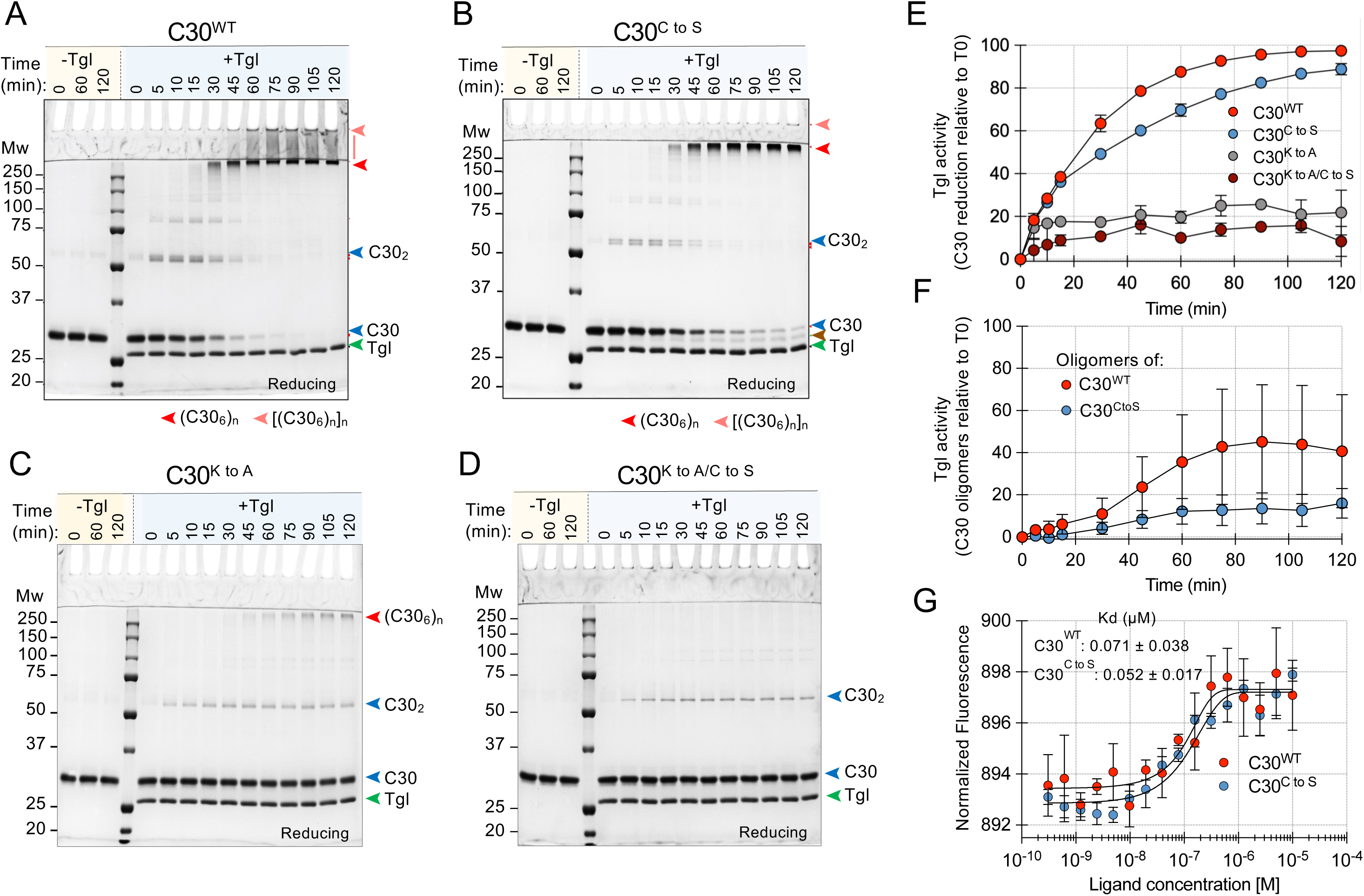
Disulfide bonds enforce the proper kinetics of C30 cross-linking by Tgl. C30^WT^ (**A**), C30^C to S^ (**B**), C30^K to A^ (**C**), and C30^K to A/C to S^ (**D**) were purified and incubated in the absence (-) or in the presence of Tgl (+). Samples were taken at the indicated time points (in min) and resolved by SDS-PAGE (10%) under reducing conditions. In **A**-**D**, the blue arrowheads show the position of C30 or C30_2_ whereas the red and pink arrowheads show the position of (C30_6_)_n_ or [(C30_6_)_n_]_n_. The green arrowhead shows the position of Tgl, and the brown arrowhead shows the position of a C30^C to S^ species that migrates faster than the monomer. Gels were stained with Coomassie. **E**: kinetics of cross-linking of C30^WT^, C30^C to S^, C30^K to A^, and C30^K to A/C to S^ by Tgl as measured by the rate of disappearance of C30. **F**: kinetics of formation of the [(C30_6_)_n_]_n_ species from C30^WT^ (red) and C30^C to S^ (blue) in the presence of Tgl. **G**: interaction of Tgl with C30^WT^ or C30^C to S^ measured by microscale thermophoresis. The estimated K_D_ for the interaction is indicated. The data in **E** to **G** are from three independent experiments.

We infer that disulfide bonds facilitate the cross-linking of C30, possibly by aligning and bringing the Lys residues within the right distance to Gln residues for efficient cross-linking by Tgl, while minimizing side reactions. We further infer that disulfide bonds are important for the formation of [(C30_6_)_n_]_n_.

### Role of the lysine residues in the cross-linking of C30

When C30^K to A^ with both the Lys177 and Lys318 residues replaced by Ala (above) was incubated with Tgl, the formation of high molecular weight species resistant to reducing SDS-PAGE was significantly diminished concomitantly with accumulation of C30_2_ (**Fig 6C**; see also **Fig 3**). Nevertheless, at levels much lower than for the WT, (C30_6_)_n_ was first detected at the stacking/resolving gel interface at 30 min of incubation, and its representation increased during the duration of the experiment (**Fig 6A** and **6C**; see below for the possible origin of this species). While the dimer also accumulated, no cross-linked products, (C30_6_)_n_ or others, were detected when C30^K to A/C to S^ was incubated with Tgl (**Fig 6D**). [(C30_6_)_n_]_n_ was not detected for either C30^K to A^ or C30^K to A/Cto S^ (**Fig 6C and D**). We infer that the K177 and K318 residues are important for the formation of high molecular weight complexes of C30 in the presence of Tgl, and that somehow their substitution by Ala synergizes with the Cys to Ser substitutions to block the formation of SDS-PAGE-resistant high molecular weight forms of C30.

To evaluate the individual contribution of the lysine residues in the cross-linking of C30, we purified forms of C30 with Lys177 or Lys318 substituted by Ala (K177A or K318A) and assessed their cross-linking by Tgl in comparison with C30^WT^ and with C30^K to A^, using reducing SDS-PAGE. We found that formation of both (C30_6_)_n_ and [(C30_6_)_n_]_n_ was reduced for either variant but slightly more for the K177A variant; also, C30_2_ was detected throughout the experiment, up to 120 min, whereas in the case of C30^WT^ it essentially disappeared from minute 75 onwards (**Fig 6C** and **6D**; **Fig 7A**-**7D**). Intermediate forms (C30, C30_2_, C30_3_…) were more easily detected for C30^K177A^ and fainter for C30^K318A^ suggesting that cross-linking is slower in the absence of K177 (**Fig 7B** and **7C**). In fact, while both K177 and K318 contribute for the cross-linking of C30 by Tgl, the enzyme cross-linked C30^K318A^ at the same rate as the WT whereas the kinetics of the reaction was slower for C30^K177A^ suggesting a more important role for this residue (**Fig 7E**). Importantly, bands at the position of (C30_6_)_n_ are still detected upon incubation of the K177A/K318A variant (no Lys residues), with Tgl (**Fig 7D**). These forms, also detected in the experiment documented in Figure 6, were absent when the K177A/K318A and Cys to Ser substitutions were combined in C30^K to A/C to S^ (**Fig 6**, compare panels **C** and **D**). This suggests that some Tgl-dependent cross-linking still occurs with the K177A/K318A variant but not with the C30^K to A/C to S^ variant. Several substitutes for lysine as the acyl-acceptor substrate have been described, including the short chain alkyl-based amino acid glycine, as well as the esterified α-amino acids Thr, Ser, Cys and Trp (38,39). Since the Lys-independent cross-linking is abolished in the K to A/C to S variant, we speculate that a modified Cys serves as an acyl-acceptor substrate in the absence of K177 and K318 (C30^K to A^). If so, how the Cys residues in C30 undergo esterification and why Ser does not support cross-linking is presently not known.

**Fig 7.**
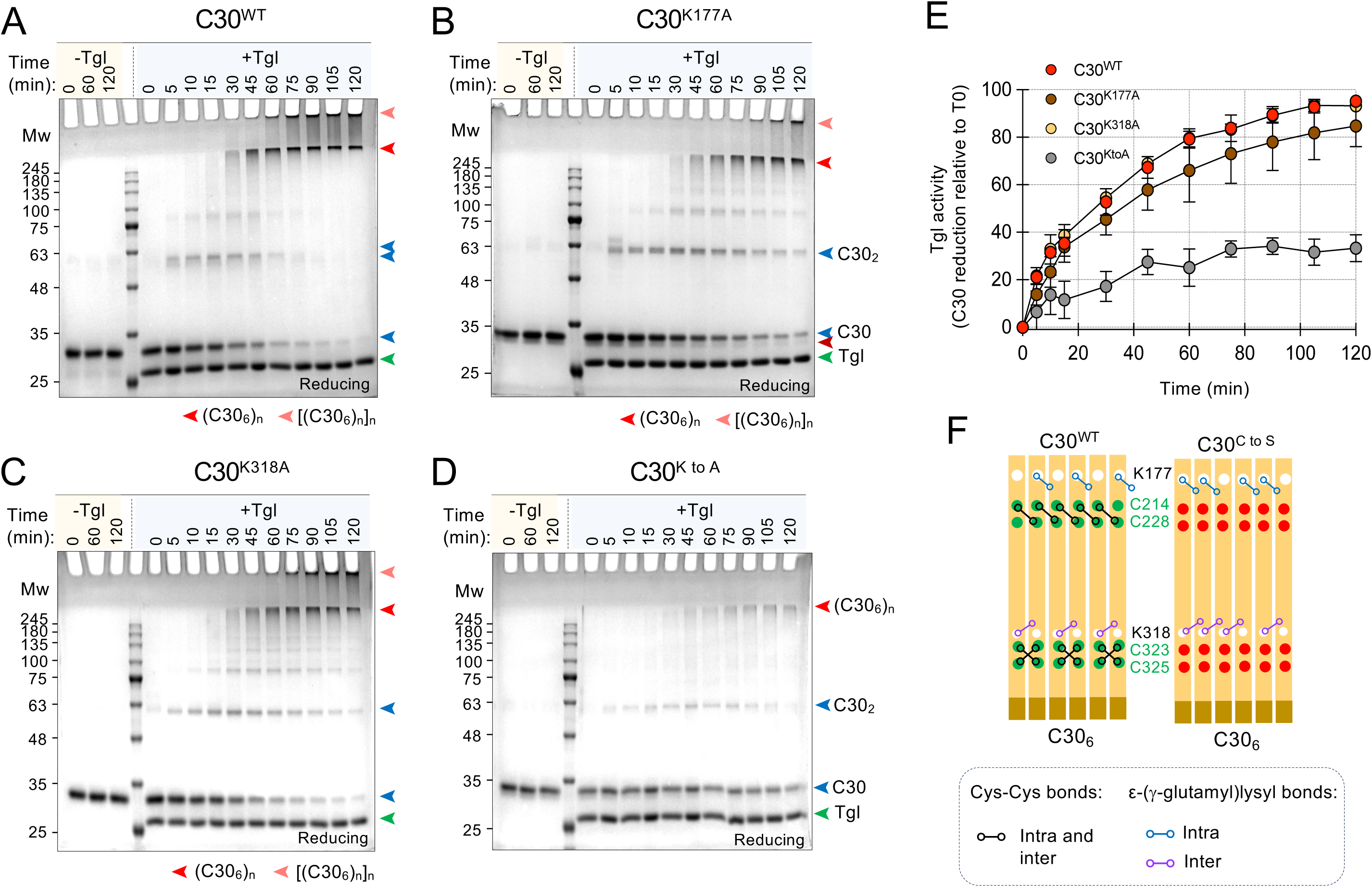
Importance of the lysine residues in the cross-linking of C30. C30^WT^ (**A**) and variants K177A (**B**), K318A (**C**), and K177A/K318A (K to A for simplicity) (**D**) were purified and incubated alone (“-“) or in the presence of Tgl (“+”). Samples were taken at the indicated times (in min) and resolved by SDS-PAGE (10%) under reducing conditions. The gels were stained with Coomassie. In **A**, **B**, **C** and **D** panels, the blue arrowheads show the position of C30 and C30_2_; the red and the faint red arrowheads show the position of HMW forms. The green arrowhead shows the position of Tgl. **E**: kinetics of cross-linking of each variant was measured by the rate of disappearance of the C30 monomer. The graph shows the data obtained with three independently purified C30 and cross-linking assays. **F**: model for the role of K177 and K318 in formation of the C30 cross-linked oligomers. In C30^WT^, K177 is involved in intramolecular (intra-dimer) cross-links, whereas K318 is thought to be involved in intermolecular (between pairs of dimers) cross-links. The panel also represents the altered pattern of C30^C to S^ cross-linking.

Formation of the cross-linked, SDS-PAGE-resistant (C30_6_)_n_ and [(C30_6_)_n_]_n_ complexes only requires the presence of either K177 or K318 establishing intra- (within C30 dimers) or intermolecular cross-links (between adjacent dimers) with one of the various Gln residues in an adjacent C30 molecule. In other words, C30 molecules that are not cross-linked via one lysine residue are cross-linked via the other (**Fig 7F**). As we described above, the N-terminal Cys residues are more important for the formation/stability of high molecular weight complexes of C30 (**Fig 4B**). Since K177 has a slightly more important role in the accumulation of (C30_6_)_n_ and [(C30_6_)_n_]_n_ than K318, we suspect that K177 may only form intermolecular cross-links whereas K318 may only make intramolecular cross-links (**Fig 7F**).

### C30 complexes adopt a globular, slightly elongated conformation

Having shown that disulfide bonds are important for proper oligomerization and the subsequent cross-linking of C30 by Tgl, we next wanted to gain insight into the structural transitions, if any, resulting from the cross-linking of the complexes. We used size-exclusion chromatography coupled with solution small-angle X-ray scattering (SEC-SAXS) to analyze C30^WT^ before and after cross-linking by Tgl (see **Fig 8A**). The SEC-SAXS chromatograms of C30 show a single symmetric peak suggesting the presence of a dominant particle at least in the area shaded in blue (**Fig 8A**); in fact, the radius of gyration (Rg) is not constant across the peak indicating the presence of higher order structures (**Fig 8A**, red dots). The distribution of the Rg values clearly indicates that the co-existence of species of different sizes (**Fig 8A**, inset). We then incubated C30 with Tgl under the same conditions used for the cross-linking assays described above (see for instance **Fig 5A** and **5B**). The chromatogram for C30 after incubation with Tgl shows a bimodal pattern, with a new peak emerging to the left (**Fig 8A***, p* in the bottom panel). This indicates formation of a species that is larger than and/or has an altered shape relatively to the main one (**Fig 8A**, arrow). The distribution of the Rg values clearly shows that *p* is populated by larger species (**Fig 8A**, inset). While the dominant peak may correspond to the species detected in the cross-linking assays at the stacking/resolving gel interface, or (C30_6_)_n_, the smaller peak may correspond to the larger species that remains at the well, or [(C30_6_)_n_]_n_ (see for example **Fig 5A**, bottom panel). Unfortunately, *p* does not show constant Rg values, precluding its analysis. In any event, the constant Rg values on the right side of the dominant peak obtained for C30 and for C30 + Tgl, indicate a monodisperse region (**Fig 8A**).

**Fig 8.**
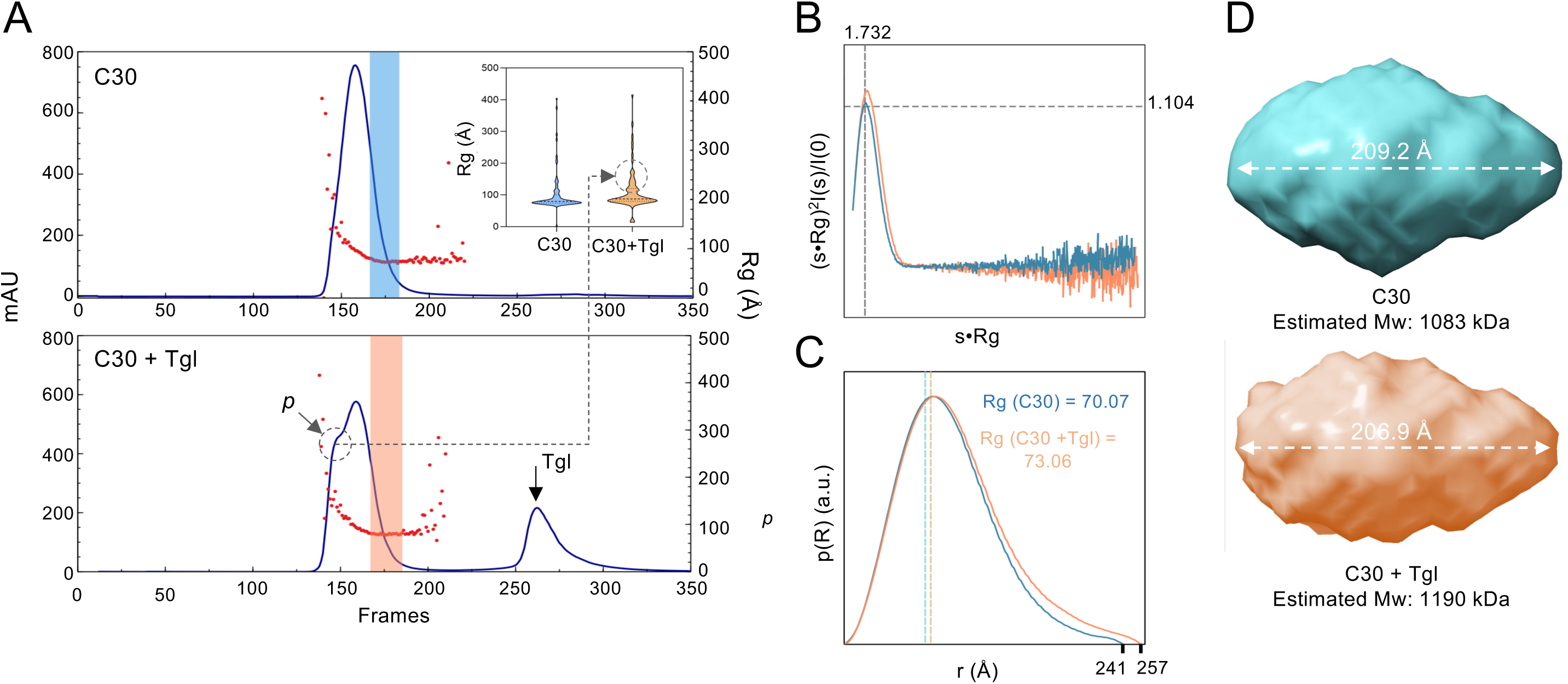
Models of C30^WT^ oligomers. **A:** SEC-SAXS chromatograms of C30 (top) and C30 after incubation with Tgl (C30 + Tgl; bottom). The UV absorbance at 280 nm is shown across the elution frames; the corresponding radius of gyration (Rg) are shown as red dots. The inset shows violin plots of Rg distributions across each peak; for the C30 + Tgl sample, a dotted line links peak *p* to the Rg distribution plot. The position of Tgl is indicated by a black arrow. **B**: Kratky plots of SAXS profiles for C30 (blue) and C30 + Tgl (green), averaged from frames with constant Rg values, shown alongside the distribution of their P(r) versus (same color). **C**: Normalized pair-distance distribution functions, P(r), derived from the experimental SEC-SAXS data for C30 (blue) and C30 + Tgl (green). **D**: Low-resolution *ab initio* reconstructions (DAMFILT of 20 independent runs) of C30 (light blue) and C30 + Tgl (green) show elliptical, globular particles with a dense core, suggesting a compact nature of the assemblies. The estimated Mw of the complexes is indicated.

The C30 elution peak corresponds to particles with an average Rg of 103.3 ± 62 Å; this value increases to 165.8 ± 62 Å upon incubation with Tgl (**Fig 8A**). Kratky plots of the SAXS profiles for C30 and C30 following incubation with Tgl, averaged from frames with stable Rg values (see above), suggests a globular shape for the eluting species (**Fig 8B**). The normalized pair-distance distribution functions, P(r), derived from the experimental data for C30 and C30 with Tgl, further support the existence of relatively small differences in size and shape (**Fig 8C**). Finally, the *ab initio* reconstructions of C30 and C30 after incubation with Tgl show elliptical, globular particles with a dense core, consistent with oligomers with a compact nature (**Fig 8D**).

Importantly, the Mw estimated, for C30 (1083 kDa) is in the range of the size of uncross-linked C30 complexes, termed (C30_6_)_n_, estimated by BN-PAGE (**Fig 5B**). Thus, (C30_6_)_n_ may correspond at least to an hexamer of hexamers (*i.e*., n = 6) which forms independently of Tgl and is subsequently cross-linked by the enzyme. Following cross-linking with Tgl, the overall shape of the complexes is maintained but the size increases slightly, to 1190 kDa (**Fig 8C**). The estimated size for the complexes (1083 kDa uncross-linked, 1190 kDa cross-linked) are in the range of the sizes estimated by BN-PAGE (**Fig 3C**). It seems possible that at least some of C30 complexes, those in the “Rg-stable” region of the chromatogram, remain largely unchanged following cross-linking by Tgl in line with the idea that the complexes are “spotwelded”.

### Tgl reduces the extractability of SafA^FL^ from spores

To examine the impact of the C30 K to A and C to S substitutions in the assembly and functionality of the spore surface layers, we first examined the protein composition of the coat/crust layers. Spores were purified from cultures of the various strains and the spore coat and crust proteins were extracted. The proteins were resolved by SDS-PAGE and the gels subject to immunoblot analysis with an anti-SafA antibody. In these experiments, our reference strain was a *ΔsafA* mutant complemented in trans, at *amyE*, with the *safA*^WT^ allele (**Fig 9A**); for direct comparison, the various K to A and C to S alleles (“cross-linking mutants”) were introduced at the *amyE* locus of the *ΔsafA* mutant. Several observations characterized our group of strains: *i*) the collection of coat/crusts proteins extracted from spores of the reference strain did not differ significantly from the “normal” WT, as judged by Coomassie staining (**Fig 9A**, top panel); *ii*) several proteins were missing or their extractability reduced from *ΔsafA* spores, as compared to the WT, including the CotG protein, as reported before (**Fig 9A**, red arrowheads) (14,17); *iii*) SafA^FL^ and C30 (blue arrowheads), GerQ and YeeK (back arrowheads) were more extractable from *Δtgl* spores, as expected (**Fig 9A**, top) (14,17).

**Fig 9.**
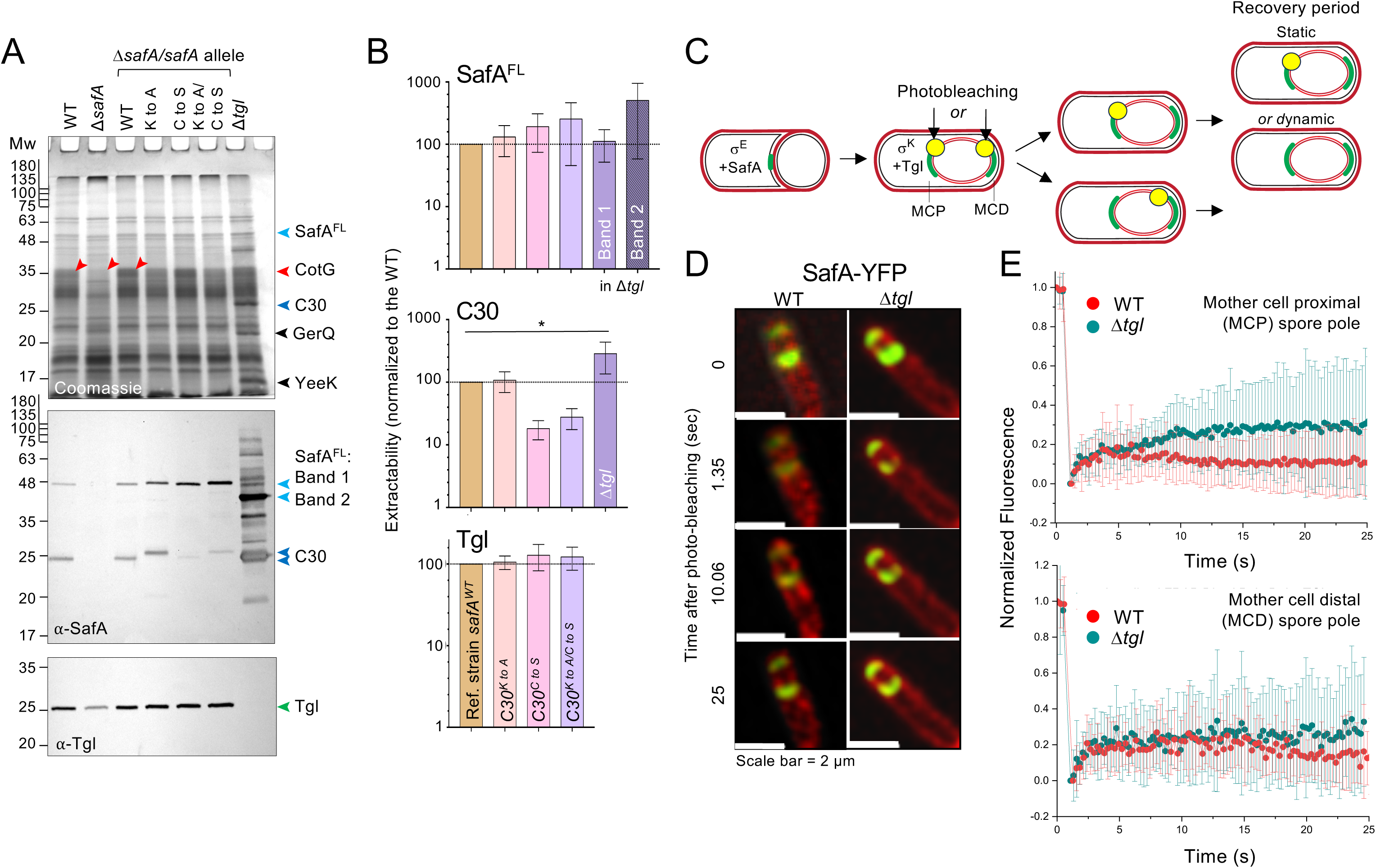
Tgl controls the extractability and dynamics of SafA^FL^/C30 complexes. **A**: Spores of the following strains were purified, the coat proteins extracted and resolved by SDS-PAGE (15%): the WT, a *safA* in-frame deletion mutant (*ΔsafA*) and a *tgl* deletion mutant (*Δtgl*); other strains have a *ΔsafA* deletion and are complemented in trans, at the non-essential *amyE* locus with the WT, *safA*^C to S^, *safA*^K to A^ and *safA^K to A/C to S^* alleles. The gels were stained with Coomassie (top panel) or immunoblotted with anti-SafA (middle) or anti-Tgl antibodies (bottom). The arrowheads show the position of the following proteins: SafA^FL^, bands 1 and 2, light blue; C30, dark blue; GerQ and Yeek, black; CotG, red; Tgl, green. Note that the K to A substitutions retard the migration of SafA^FL^ or C30. **B:** quantification of the levels of the various proteins extracted from spores of the *ΔsafA* mutant complemented with the indicated *safA* alleles at *amyE*. The data are from three experiments using independently prepared spore suspensions and shown normalized to the WT. Note that the quantification of bands 1 and 2, not seen in the WT, is relative to SafA^FL^. For the statistical analysis we used a one-way ANOVA followed by Dunnett’s multiple comparison test (⍺ = 0.05). Asterisks indicate statistically significant differences (*p* < 0.05), with * denoting values <0.05 relative to the WT. **C:** Represents cells at the time when SafA begins to be synthesized under σ^E^ control and localizes to the spore surface, and cells after engulfment completion, when Tgl is produced under the control of α^K^. At this stage SafA forms caps at the MCP and MCD forespore poles. For both poles, the edge of the cap was photobleached and the signal recorded over time. The recovery period is only illustrated for the MCP pole; the protein is dynamic, if the signal recovers, or otherwise static. **D**: Cells were collected 5 hours after the onset of sporulation and imaged by confocal microscopy. The signal from SafA-YFP was photobleached at the MCP forespore pole in the WT and in *Δtgl* mutant. **E**: The signal in the photobleached areas was then monitored for up to 25 seconds and plotted with the standard deviation for the mother cell proximal (top) and distal (bottom) forespore poles.

The immunoblot analysis revealed that in comparison with the reference strain, SafA^FL^ was slightly more abundant or more extractable from spores of the SafA^K to A^, SafA^C to S^ and SafA^K to A/C to S^ mutants (**Fig 9A**, mid panel; quantification in **9B**, top panel; note that SafA^FL^ or C30 bearing substitutions of the Lys residues to Ala migrate slightly slower than other forms of the protein). Thus, consistent with the idea that Tgl cements SafA^FL^/C30 complexes at the cortex/inner coat interface, the SafA^FL^ variants with the substitutions that impair cross-linking are more extractable from mature spores.

Interestingly, C30 was less abundant or less extractable from the C to S or K to A/C to S mutants but not from the K to A mutant (**Fig 9A**, mid panel; **Fig 9B**, top panel). Possibly, the disulfide bonds formed between C30 or between C30 and the C30 domain of SafA^FL^ are important for their interaction, which in turn is required for the localization of C30 to the coat, explaining its reduced abundance/extractability (23).

In *Δtgl* spores no band was detected with the migration of SafA^FL^ in extracts from the reference strain; there were, however, several species below including band 1 (**Fig 9A**, middle panel) and a prominent band at around 40 kDa (band 2; quantification relative to SafA^FL^ is shown in **Fig 9B**). Species below the position of SafA^FL^ in the reference strain (such as band 1 and band 2) suggest that SafA^FL^ is subject to proteolysis in the absence of Tgl, whereas the species above the position of SafA^FL^ suggest Tgl-independent cross-linking (**Fig 9A**, middle panel). Overall, more SafA^FL^ and C30 are extracted from *Δtgl* spores than from spores of the K to A/C to S strain (**Fig 9A**, middle panel) suggesting Tgl-dependent cross-linking to other proteins. These are likely to be SafA^FL^-dependent, such as YeeK and GerQ, which are also Tgl substrates (**Fig. 1C**). Importantly, with the exception of the *ΔsafA* mutant (SafA recruits Tgl; (25) and the *Δtgl* control, the level of Tgl in spores of the K to A and C to S mutants was comparable (**Fig 9A** and **9B**, bottom panel). Therefore, changes in the extractability of SafA^FL^ or C30 from spores of the various “cross-linking mutants” are not due to differences in the amount of Tgl that associates with the spores.

A *ΔsafA* mutant forms spores with compromised coat layers that are permeable to and sensitive to lysozyme; these spores are also heat sensitive, which indicates a defective spore cortex peptidoglycan (14,15). Spores of a mutant expressing *C30^K to A/C to S^* showed a small but reproducible decrease in heat and lysozyme resistance (**S9A_Fig**), suggesting that proper cross-linking of SafA^FL^ and C30 is important for proper assembly of the spore coat layers. None of the mutants showed impaired spore germination as compared with the reference strain (*ΔsafA* with *safA* ^WT^ at the non-essential *amyE* locus; **S9B_Fig** and **S9C_Fig**).

### Tgl immobilizes SafA^FL^ and C30 at the spore surface

The increased extractability of SafA^FL^ and C30 from spores of a *Δtgl* mutant suggested to us that the Tgl-dependent cross-linking of C30 could immobilize the protein within the coat. The *safA* gene is expressed early in development, under the control of α^E^. A functional SafA-YFP (note that because of alternative translation initiation C30-YFP is also produced, *in vivo*, from the same construct (15) localizes at the mother cell proximal (MCP) forespore pole, at the onset of engulfment and forms caps at both the forespore poles soon after engulfment completion when σ^K^ is activated and Tgl is produced (**Fig 9C**; see also **Fig 1A**) (40). We used fluorescence recovery after photobleaching (FRAP) to examine the dynamics of SafA-YFP at the mother cell proximal (MCP) and distal (MCD) forespore poles in the WT and a *Δtgl* mutant. We photobleached an area of the forespore membranes at the MCP and MCD poles, in strains producing SafA-YFP, and imaged the cells over time, on a confocal microscope, to monitor FRAP (**Fig 9C** and **9D**, top panel). Fluorescence recovery would indicate incorporation SafA-YFP or the exchange of SafA-YFP molecules at the site of photobleaching and thus a dynamic behavior whereas no recovery would indicate formation of static complexes (**Fig 9C**). In the *Δtgl* mutant, but not in the WT, about 20% of the photobleaching levels of SafA-YFP recovered by 15 sec, but only at the MCP of the forespore (**Fig 9E**). Although we do not presently know why the two spore poles differ, we conclude that Tgl immobilizes SafA^FL^/C30 complexes at the spore surface, consistent with the increased extractability of SafA^FL^ variants K to A, C to S and K to A/C to S from *Δtgl* spores (**Fig 9A**, mid panel; **9B**, top panel).

## DISCUSSION

The intrinsically disordered C30 domain acts as a dual Hub, shared between two proteins, SafA^FL^ and C30, and is responsible for recruiting inner coat proteins. This Hub-like role is particularly evident in mutants expressing C30 in the absence of the full-length protein: without localization signals, C30 remains in the mother cell cytoplasm, where it serves as a strong attractor for the entire coat (25). Within the IDP class, C30 belongs to the “globule” category and likely harbors multiple self-interaction hotspots (30,32) (**Fig 2D**), which facilitate its self-assembly into large complexes. A useful parallel can be drawn with amelogenin, another IDP that self-assembles into diverse supramolecular structures involved in dental enamel formation. Amelogenin transitions from monomers (at pH < 3.5) to ∼8-mer oligomers (at pH 6.8), to nanospheres of 20–100 monomers (at pH > 7), and can form chains of nanospheres or nano- and micro-ribbons under specific conditions (33–35). Similarly, the oligomerization of C30 appears critical for inner coat assembly (**Fig 10**).

**Fig 10.**
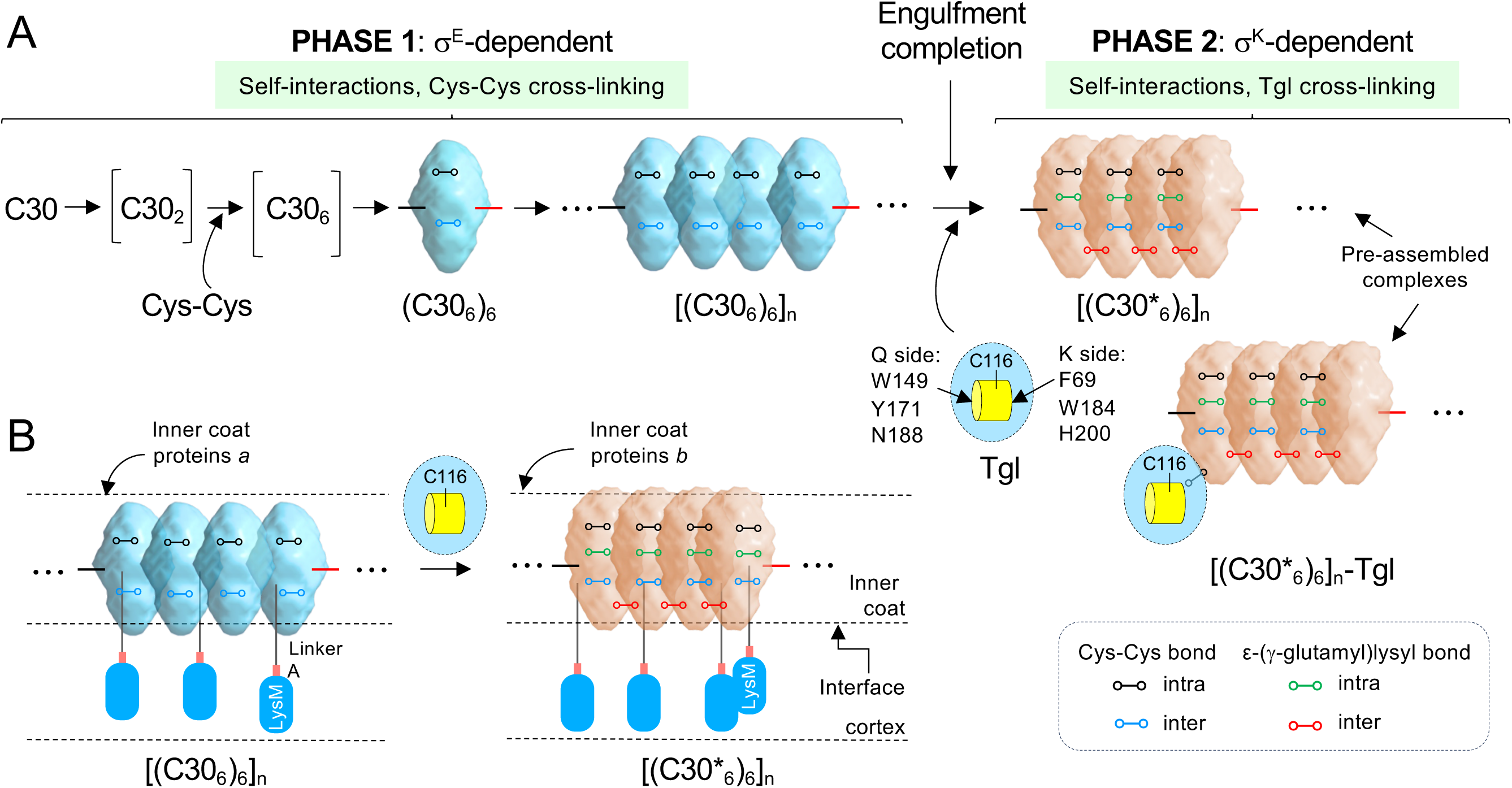
Assembly of SafA is biphasic and relies on a hierarchical cross-linking cascade. **A.** Prior to engulfment completion, SafA^FL^ (not shown for simplicity) and C30 are produced and self-assemble into oligomers that undergo disulfide cross-linking. Following engulfment completion, activation of σ^K^ leads to production of Tgl, which is recruited to the spore surface via pre-assembled SafA^FL^/C30 complexes. Tgl then *“*spotwelds” these complexes resulting in their immobilization and reduced extractability. Tgl itself may also become cross-linked to the SafA^FL^/C30 complexes. **B.** C30 relies on the interaction with SafA^FL^ for localization to the cortex/inner coat interface. Each C30 hexamer may incorporate one or more SafA^FL^ molecules, anchoring the complex to the cortex. SafA^FL^/C30 complexes may recruit different inner coat proteins (*a*, *b*) before or after Tgl-mediated cross-linking (asterisks). The complexes are not drawn to scale and the CAM_6_ motif is omitted for simplicity.

The assembly pathway of the SafA^FL^/C30 dual Hub is biphasic. Phase 1, from the onset of engulfment to completion of the process, is dominated by self-assembly and disulfide bond formation; phase 2 begins at engulfment completion when σ^K^ is activated, and is marked by the Tgl-dependent cross-linking of the pre-formed complexes. Assembly of C30 most likely begins with hydrophobic, DDM-labile, interactions and with the parallel orientation of dimers (**Fig 3C** and **S7_Fig**). It proceeds with the cross-linking of the complexes through disulfide bonds in a criss-cross pattern involving two pairs of N-terminal Cys residues, Cys214 and Cys228, and two pairs of C-terminal Cys residues, Cys323 and Cys325. In the absence of the N-terminal Cys residues, the C-terminal disulfide bonds are sufficient for the formation of a cross-linked tetramer, possibly a dimer of dimers, but oligomers of higher order do not accumulate as assessed by BN-PAGE. In contrast, in the absence of the C-terminal disulfide bonds, C30 still forms large complexes, at least (C30_6_)_n_ (**Fig 3B** and **3C**). Nevertheless, the C-terminal bonds have a role in the stabilisation of C30 oligomers because in the absence of all of the Cys residues, either by treating C30^WT^ with DTT or by analysing C30^C to S^, only an uncross-linked C30 dimer is detected by BN-PAGE and the complexes found by SEC are smaller (≈ 762 kDa) than for the WT (≥1235 kDa) (**Fig 3C** and **3D**). The dimers and tetramers observed in the absence of Cys residues suggests that formation of the oligomers drives disulfide bond formation and not vice versa (41). Past the dimer/tetramer stage, however, it is unclear whether disulfide bond formation stabilizes and/or are the driver for the assembly of higher order oligomers.

Formation of the Cys214-Cys228 and Cys323-325 bonds requires the proximity and correct orientation of the intervening side chains (41). Residues Cys214 and Cys228 are predicted to be within a hotspot region of C30 (residues 191-316; **Fig 2E**); the Cys323/Cys325 pair, however, is at the C-terminal fringe of this region. The apparently more important role of the N-terminal pair of Cys residues in self-assembly thus seems to match the propensity of the vicinity of Cys214 and Cys218 to self-interact (**Fig 2E)**. Self-interaction may bring Cys214 and Cys228 in close proximity and with the correct orientation for disulfide bond formation; this, in turn may generate a signal that propagates along the molecule facilitating the disulfide cross-linking of the C-terminal pair of Cys residues. In this model, the Cys-Cys bonds in the SafA^FL^/C30 double Hub are primarily structural, but those at the N-terminal end of the C30 domain further have an allosteric function (42). Interestingly, NMR studies indicate that the self-assembly of amelogenin begins with the parallel alignment of monomers and starts with intermolecular interactions at the N-terminal TRAP region with the later involvement, through weaker interactions, of a small region near the C-terminus (33–35) (**S2_Fig**).

Importantly, since K177 and K318 are also at the fringes of the hotspot region, the disulfide bonds may orient parallel C30 molecules in a way that brings the lysine residues in the proximity of an acyl-donor Gln for their cross-linking by Tgl. The Cys-Cys bonds are thus at the basis of a hierarchical cascade of protein cross-linking that intervenes in the assembly of the C30 Hub. In their absence, C30^C to S^ oligomers are still cross-linked but the kinetics of the reaction is slower and the large [(C30_6_)_n_]_n_ species are not formed (**Fig 6**). The SAXS experiments revealed that complexes of C30 with about 1083 kDa are cross-linked by Tgl and remain largely unchanged (**Fig 8**). This is in line with the idea that the enzyme exercises a “spotwelding” activity, eventually cross-linking all of the subunits in a pre-formed complex (**Fig 6A**, **Fig 7A** and **S8_Fig**). This spot-welding activity is also seen in the blood coagulation cascade, for instance, where the Factor XIII transglutaminase intervenes, and may be a general attribute of transglutaminases (43). In the process, Tgl itself becomes part of the complexes which *in vivo* may be a way to shut down activity (**S3_Fig** and **S4_Fig**) (25). In spite of having several Lys residues, the LysM domain and the linker are not substrates for the enzyme *in vitro* (**S1A_Fig** and **S5_Fig**); thus, the C30 domain is likely to be the only part of SafA^FL^ that is cross-linked by Tgl. The increased extractability of SafA^FL^ and C30 from *Δtgl* spores is consistent with the cross-linking of the protein to itself or to C30 (**Fig 9A**). In sporulating cells, soon after engulfment completion and in the absence of Tgl, SafA^FL^-/C30-YFP molecules are more mobile at the MCP forespore pole but remain essentially static at the MCD pole (**Fig 9D** and **9E**). One possibility is that encasement by both SafA^FL^/C30 and Tgl normally initiates at and proceeds from the MCP pole so that complexes become immobilized first at this pole. In any event, these observations are in keeping with the idea that Tgl cements the cortex/inner coat interface.

Hubs may be static (or party) or dynamic (date) (3,44); static Hubs bind several partners simultaneously at different locations whereas a dynamic Hub tends to bind short linear motifs, or SLiMs, in intrinsically disordered regions of otherwise globular proteins and has several partners competing for the same site (3). SLiMs may only provide weak interactions between the Hub and partner proteins but this may be sufficient for the Tgl-C30 enzyme-Hub interaction and Tgl eventually auto-cross-links to the Hub. Stronger binding may involve oligomerization of at least one side of the Hub-client protein interface. The strong self-interactions of the C30 domain and the formation of large oligomers such as (C30_6_)_n_ and [(C30_6_)_n_]_n_, of which C30_6_ may be the structural repeating unit, suggests a model in which the Hub forms a continuous shell around the forespore, cross-linked by Tgl. Docking motifs are ligand-binding motifs that act as recruitment sites for client proteins (3). While the C30 client proteins may bind to the same docking site in C30, *i.e*., C30 acting as a dynamic Hub, the abundance and oligomerization of the domain may render recruitment essentially non-competitive. An alternative view is that the SafA^FL^/C30 Hub combines features of both static and dynamic Hubs, both during the σ^E^- and α^K^-dependent phases (**Fig 9** and **Fig 10**).

Recruitment of the protein-modifying Tgl by SafA^FL^/C30 illustrates binding to docking motifs in the Hub that become modified by formation of a ε-(γ)-glutamyl-lysine cross-link and therefore unlikely to be competed for by other partners (3). Tgl first engages an acyl donor Gln to form an acyl-enzyme intermediate, and then engages a motifs comprising an acyl-acceptor Lys residue to complete the cross-linking reaction (25). This sequence directs Tgl to its substrates but may not be stringent enough to explain enzyme specificity (23,25) (**Fig 1D**). Modification enzymes often have additional binding sites for docking motifs in the Hubs in a surface distinct from the catalytic centre, that complement the enzyme-modification site on the Hub (3). The substrate-driven localization of Tgl results in the spatial and temporal co-localization with its substrates which in turn ensures that the correct substrates are cross-linked and off-site reactions are minimized. A second, substrate-independent pathway, however, may involve a binding surface different from the active site region in Tgl and a different docking site in the Hub (25). This interface has not yet been identified.

Immunogold labelling showed that the Hub, and also Tgl, are located at the cortex/inner coat interface (23). That SafA relies on an interaction with SpoIVA for targeting and with SpoVID for encasement, suggests that the Hub has an internal localization also in sporulating cells. Such a localization is further supported by recent cryo-electron tomography (cryo-FIBM/ET) studies of sporangia suggesting that a layer that may contain SafA is located at about 50 nm from the OFM (45). In one model SafA^FL^/C30 exists as two sub-populations, one closer to the OFM, the other in association with the inner coat cross-linked by Tgl in the two locations (23). If SafA^FL^ is attached to the underlying cortex via its LysM domain, then the length of the linker region may be important to define the distance from the cortex where the C30 domains resides and hence the site of inner coat assembly. To resolve the molecular details on how the Hub complexes and Tgl organize the cortex/inner coat interface to pattern the assembly of the inner coat proteins is a key objective for future work.

## Material and Methods

### Bacterial strains, media and general techniques

The *B. subtilis* strains are all congenic derivatives of strain MB24. The *Escherichia coli* strain DH5⍺ (Bethesda Research laboratories) was used for molecular cloning. The *E. coli* strain BL21(DE3) (Novagen) was used for recombinant proteins expression. Luria-Bertani medium was routinely used for growth and maintenance of *E. coli* strains. Solid media were prepared by adding 1.6% (w/v) agar to liquid media. All PCR products or plasmids constructed were verified by sequencing. All strains and their relevant properties, primers and plasmids used in this study are listed in **S3_Table**, **S4_Table** and **S5_Table**, respectively.

### SDS-PAGE and immunoblotting

Proteins were resolved by SDS-PAGE (different concentrations were used as indicated in the sections below or in the Supplemental Information). The gels were stained with Coomassie or subject to immunoblot analysis with an anti-SafA or anti-Tgl antibodies at a dilution of 25000 and 1:15000 respectively; a secondary anti-rabbit peroxidase-conjugated antibody was used at a concentration of 1:10000. The immunoblots were developed using the SuperSignal West Pico Chemiluminescent Substrate kit according to the manufactureŕs instructions (Thermo Scientific) and imaged using the *i*Bright FL1500 Imaging System (Invitrogen).

### Blue native gel electrophoresis

Mono-dimensional blue native polyacrylamide gel electrophoresis (1D BN-PAGE) was used to resolve the proteins in their native oligomeric states. The gels consisted of a stacking gel (5% acrylamide) and a resolving gel (gradient of 5-15% acrylamide).Before electrophoresis, proteins samples as well as protein marker were incubated for 30 min at 37°C with BN-PAGE loading buffer (0.1% Coomassie blue, 20mM Tris-HCl, pH 7.6) and 0.05%-dodecyl-β-D-maltoside (DDM). Electrophoresis was carried out at 4 mA for 30 min at RT, then for 1h30 at 4°C at 6mA. The anode buffer consisted of 50 mM Bis-Tris, pH 7, and the cathode buffer was 15 mM Bis-Tris, 50 mM tricine, supplemented with 0.02% Coomassie Brilliant Blue G-250 (Sigma). Gels were stained with Coomassie. NativeMark™ Unstained Protein Standard (Invitrogen™) was used as a molecular weight (Mw) marker.

### Size exclusion chromatography

For Size Exclusion Chromatography (SEC), a Sephacryl S-300 HR 120 mL column (Cytiva) was equilibrated with SEC buffer (0.1M Tris-HCl, 0.15 M NaCl, pH.8); 1 ml of Ni^2+^-affinity purified C30^WT^ or C30^C to S^ proteins (1 mg/ml, unless stated otherwise), either subjected to Tgl cross-linking (as below) or un-cross-linked, were injected and the columns was run at a flow of 1 mL/min at 4°C. The column was calibrated with a Gel Filtration Standard (Bio-Rad) with the following composition: bovine thyroglobulin, 670 kDa; bovine γ-globulin, 158 kDa; chicken ovalbumin, 44 kDa; horse myoglobin, 17 kDa; vitamin B12, 1.35 kDa.

### Cross-linking assays

Each of the C30 variants (12 µM) was incubated with Tgl (4µM) in a volume of 100 µl 0.1M Tris-HCl, pH 8.0, for up to 2h at 37°C; 5 µl samples were collected at various times and incubated for 1 min at 100°C in 10x loading buffer either reducing [Tris-HCl 62.5 mM pH6.8, SDS 2% (w/v), glycerol 25% (v/v), 0.5mM DTT, 2-mercaptoethanol 0.5% (v/v) bromophenol blue 0.02% (w/v)] or non-reducing (DTT and 2-mercaptoethanol omitted). Samples were resolved by SDS-PAGE (10% gels). To assess the activity of Tgl we measured the loss of C30^WT^ or its variants over time on Coomassie-stained gels, using the iBright analysis software. The values obtained were normalized according to the following formula (where t represents the different time points, in minutes, during a cross-linking reaction) (25):

Protein Cross-linking_norm_ = (1-(Protein at time t)/(Protein at time 0))*100.

The crosslinking reactions of SafA^FL^ and its N-terminal domain, SafA^N21^, were performed similarly to C30 but at different concentrations. Briefly, in a first series of assays 4 μM Tgl was incubated with increasing concentrations of either SafA^FL^ or SafA^N21^. In a second series, SafA^FL^ at 12 µM was incubated with increasing concentrations of Tgl. Samples of 5 μL were collected at the indicated time points, heated for 1 min at 100°C in 10X loading buffer and then resolved by SDS-PAGE (10% gels) in reducing conditions. Tgl activity was quantified as described above. The crosslinking reactions for SEC analysis experiments were performed in 1ml volumes of 0.1M Tris-HCl, 0,15M NaCl, pH 8.0 in 2ml *Eppendorf* tubes, maintaining the 1:3 enzyme to substrate ratio used in the PAGE resolved reactions.

### Mass Spectrometry

For the LC-MS analysis of disulfide bonds, the protein sample was denatured with 0.1% RapidGest (Waters) and digested in solution with chymotrypsin overnight, at a 1:50 enzyme:protein ratio, without prior reduction and alkylation. The sample was then acidified with formic acid and centrifuged 15000 x *g* for 15 min to remove RapidGest. The peptide mixture was analysed by LC-MS using a ZenoTOF7600 instrument (Sciex) coupled to the Waters M-class microLC. External calibration was performed using ESI calibration solution (Sciex). Peptides were separated through reversed-phase chromatography (RP-LC) using an Kinetex XB-C18 column (100 Å, 2.6 μm, 150 x 0.3 mm, Phenomenex), with a column temperature of 40 °C and a flow rate of 5 μL/min. The mobile phase consisted of Water with 0.1% FA (Solvent A) and Acetonitrile with 0.1% FA (Solvent B). The gradient was as follows: 0-2 min, 2% B; 2-32 min, 2-40% B; 32-33 min, 40-90% B; 33-36 min, 90% B; 36-37 min, 90-2% B; 37-42 min, 2% B. The mass spectrometry (MS) was set to a data-dependent MSMS method. The source parameters were set as follows: 25 GS1, 10 GS2, 35 CUR, 4.5 keV ISVF and 100 ⁰C IHT. An information dependent acquisition (IDA) method was set with a TOF-MS survey scan of 350-2000 m/z with 75 msec accumulation time. The 20 most intense precursors were selected for subsequent fragmentation and the MS/MS were acquired for 35 msec and Zeno Trapping ON with 1E5 threshold. The selection criteria for parent ions included a charge state between +2 and +8, and counts above a minimum threshold of 100 per second. Ions were excluded from further MS/MS analysis for 5 s. Fragmentation was performed using rolling collision energy with a collision energy spread of 3. The acquired data were processed using Sciex OS 3.1 and further analyzed with BioPharmaView (version 3.0, Sciex; https://sciex.com/) with the sequence of the protein of interest and using the peptide mapping workflow. The disulfide bonds Cys214-Cys228 and Cys323-Cys325 were considered.

### Fluorescence resonance energy transfer and laser scanning confocal microscopy

Overnight cultures of a WT strain and a congenic *Δtgl* mutant, both carrying an in-frame *ΔsafA* deletion and a functional *safA-yfp* fusion at the non-essential *amyE* locus, grown in LB were diluted 1:100 in pre-warmed LB. The cultures were then incubated (150 rpm, 37°C) until an OD_600_ ≈ 0.3-0.5. Cells were harvested for 10 min at 4200 x *g*, at room temperature and resuspended in Sterlini-Mandelstam (SM) medium supplemented with L-methionine (50 µg/mL) and L-tryptophan (20 µg/ml) (46). Samples of 1 ml were withdrawn 5 hours after resuspension; the cells were collected by centrifugation (2 min, 4200 x *g*), washed twice with 1 ml of phosphate-buffered saline (PBS; 137 mM NaCl, 10 mM Phosphate, 2.7 mM KCl, pH 7.4), and resuspended in 0.2 ml of PBS supplemented with the lipophilic styryl membrane dye N-(3-triethylammoniumprpyl)-4-(p-diethylaminophenyl-hexatrienyl) pyridinium dibromide (FM4-64, Molecular Probes, Invitrogen; 10 μg ml−1) (47). Cells were labelled for 2 min in the dark, collected by centrifugation (2 min, 4200 x *g*), and finally suspended in 20 μl of PBS. To visualize individual cells, 3 μL of the suspension were mounted on 1.7% agarose coated glass slides and observed on a ZEISS LSM 880 point scanning confocal microscope equipped with photomultiplier tube detectors (PMTs) and a gallium arsenide phosphide (GaAsP) detector. A prebleached image of the forespore membranes was acquired using the 514 nm laser line for excitation, with emission collected via a GaAsP detector. Bleaching was conducted using the same 514 nm laser line at maximum power for 3 seconds, again targeting the forespore membranes. The recovery of fluorescence in the bleached area was recorded under identical acquisition parameters to the prebleached image, for 25 seconds. Pixel size was adjusted to respect Nyquist sampling criteria, ensuring optimal spatial resolution for quantitative analysis. Localized photobleaching experiments were performed on representative cells from the sample populations (WT and mutant). For the mother cell proximal forespore pole (MCP), a total of 13 cells (n=13) were analysed for each strain; for the distal (MCD) forespore pole a total of 11 cells (n=11) were analysed per strain. Corrections for photobleaching, normalization, and averaging were performed using ImageJ/FIJI (version 1.54f) (48). Image analysis was conducted using ZEISS Zen 2.3 (black edition) software and ImageJ/FIJI 1.54 (48).

### Small-angle X-ray scattering

Synchrotron SEC-SAXS data were collected on the BM29 ESRF beamline (Grenoble, France). We injected 50 μL of C30 (4.7 mg/ml) or C30 (3.64 mg/ml) after incubation with Tgl (1.39 mg/ml) into a 5 ml AdvanceBio SEC 300A 4.6×300 (Agilent) size exclusion column at a flow rate of 0.4 mL min−1, collecting 1 frame per 2 seconds at 20°C, for a total of 350 frames. The SEC mobile phase consisted of 100 mM Tris at pH 8, 150 mM NaCl, at 20°C. The two-second frames were acquired using a Pilatus 2M in vacuum pixel detector (DECTRIS) at a sample-detector distance of 2.812 m and a wavelength of λ = 0.992 Å[47], covering a momentum of transfer range of 0.0085 < *s* < 0.49 Å-1. The scattering intensities from the right region of elution peaks (20-frames) were integrated, the buffer subtracted and averaged using the ScÅtter software to produce the averaged SEC-SAXS profiles. From this profile, the pair-wise distance distribution function, *P(r)*, was obtained by indirect Fourier Transform with GNOM using a momentum transfer range of 0.010 < *s* <0.35 Å-1 for C30 and 0.012 < *s* <0.33 Å-1 for C30 with Tgl. The *Rg* values were estimated by applying the Guinier approximation in the range *s* < 1.3/*Rg*. Low-resolution *ab initio* molecular envelopes were generated with the program DAMMIF (49). We built twenty independent *ab initio* reconstructions for each C30 oligomer by the “fast” annealing mode with P6 symmetry. Models were refined with DAMMIF (49), averaged, aligned, and compared using DAMAVER (50).

### Additional methods

The construction or use of *B. subtilis* and *E. coli* strains expressing various *safA* alleles at *amyE,* of plasmids for the overproduction of SafA^FL^, N21, C30^WT^ and its variants, and Tgl in *E. coli*, protein purification, microscale thermophoresis, structural bioinformatics and spore germination and resistance assays are detailed in the Supporting information (see **S1_Text**).

**S1-Text –** Supporting material and methods and supporting results and discussion.

**S1_Table –** SafA properties.

**S2_Table –** MSMS data of peptide ADHDDC^323^GC^1325^DGDHQPY carrying disulfide bond Cys323-Cys325.

**S3_Table –** Bacterial strains used in this study.

**S4_Table –** Oligonucleotide primers.

**S5_Table –** Plasmids used in this study.

## ACKNOWLEDGEMENTS

We acknowledge members of the group for critical reading of the manuscript and valuable suggestions. We also acknowledge the use of the *Bacterial Imaging Cluster* platform (www.itqb.unl.pt.bic) and *European Synchrotron Radiation Facility* (ESRF)

BM29 BioSAXS beamline.

## DATA AVAILABILITY STATEMENT

All relevant data are contained within the manuscript and its Supporting Information files.

## COMPETITIVE INTERESTS

The authors have declared that no competitive interest exist.

## AUTHOR CONTRIBUTIONS

**Conceptualization:** AOH, MS, ASP, TNC

**Data curation:** KA, CF, DQPR, DS, CO, BG, TP, MLM, GH, CT, RAG, MS, AOH

**Formal analysis:** KA, AOH, MS, TNC, ASP

**Funding acquisition:** AOH, MS, ASP, TNC

**Investigation:** KA, CF, DQPR, DS, CO, BG, TP, MLM, GH, CT, RAG, MS, AOH

**Methodology:** KA, EM, TP

**Project administration:** AOH, MS, TNC, AP

**Resources:** DQPR, MLM, GH, CT, RAG

**Supervision:** AOH, MS, TNC, AP

**Validation:** KA, CF, DQPR, DS, CO, BG, TP, MLM, GH, CT, RAG, MS, AOH

**Visualization:** DQPR, DS, MLM, GH

**Writing original draft:** KA, AOH

**Writing, reviewing and editing:** KA, MS, ASP, TNC, AOH

## SUPPORTING INFORMATION

## SUPPORTING MATERIAL AND METHODS

### Spore germination assays

Density gradient purified spores were heat-activated at 65°C for 15 minutes and then added to two different buffers to an optical density at 580 nm (OD_580_) of 0.5, in a final volume of 200 µl, in a 96 well plate. The buffers used were: Buffer 1: 50mM Tris-HCl, pH 7.5; Buffer 2: 50 mM Tris-HCl, pH 7.5, 3.3 mM L-asparagine, 5.6 mM D-Glucose, 5.6 mM D-Fructose, 10 mM KCl (14). To prevent spore clumping and adsorption to the plate wells, 0.01% Tween 20 was included in all buffers (54,55). The GFK mixture was freshly prepared before each experiment from stock solutions stored at 4°C. Each sample was analyzed in duplicate (two technical replicates). Germination was monitored using the BioTek Plate Reader Neo2 Multimode Reader by measurement of the decrease in the OD_580_ for up to 60min at 37°C. Germination kinetics data were analysed by calculating the average of each technical replicate at each point. Then, buffer-only values were subtracted from their corresponding spores-containing buffer values. Each data point of the kinetic experiment was then normalized to the fourth time point (at 4.36 min), as this is the point at which the measurements begin to stabilize or decrease, depending on the strain tested. The percentage of germination was calculated by normalizing the OD_580_ values at t = 60 min for each spore suspension to the corresponding value at t = 0 min. The resulting values were further normalized to the reference strain (the *ΔsafA* mutant with the *safA^WT^* allele at *amyE*).

**Constructs for the overproduction of SafA^FL^, SafA^N21^, C30 and C30 variants. SafA^FL^ and C30**: the constructs for the overproduction of SafA^FL^ and C30 were described before (Fernandes et al., 2019).

**C30^C to S^ and C30^K to A/C to S^**: The sequence coding for the C30 variants was custom-made synthesized by Blue Heron substituting the four Cys residues by Ser residues (C30^C to S^) or together the four Cys residues by Ser residues and the two Lys residues by Ala residues (C30^K to A/C to S^). The resulting fragments were inserted between NcoI/XhoI cloning sites of pET28a+ yielding pKA32 and pKA33 which were transformed into BL21 (DE3) originating AHEC1308 and AHEC1309, respectively.

**C to S^N-ter^:** The Quick-Change site-directed mutagenesis strategy was used to replace both Cys214 and Cys228 by Ser in C30 using the primer pair CtoS_51/65_Fw and CtoS_51/65_Rev (NB: Cys51 and Cys65 in C30 are Cys214 and Cys228 in SafA^FL^). pCF68 was used as template generating pKA17 which was then transformed into BL21 (DE3) originating AHEC1428. pKA17 carries C30^C214/C228 to S^ for simplicity the protein product was named C30^C to SN-ter^ indicating that the N-terminal Cys residues were substituted by Ser residues.

**C to S^C-ter^:** The Quick-Change site-directed mutagenesis strategy was used to replace both Cys323 and Cys325 by Ser in C30 using the primer pair CtoS_160/162_Fw and CtoS_160/162_Rev (NB. Cys160 and Cys162 in C30 are Cys323 and Cys325 in SafA^FL^). pCF68 was used as template to generate pKA18 which was then transformed into BL21 (DE3) originating AHEC1429. pKA18 carries C30^C323/C325 to S^ and for simplicity the protein product was named C30^C to SC-ter^ indicating that the C-terminal Cys residues were substituted by Ser residues.

**C30^K177A^ and C30^K318A^:** Plasmid pCF68, described before, carries the C30-encoding part of *safA*, from the M164 codon onwards, in pET28a(+) (25). Codons K177 and K318 were then individually replaced by A using pCF68 as the template and primer pairs safAK177A D/safAK177A R and safAK318A D /safA K318 R, respectively. This created plasmids pCF138 (K318A) and pCF146 (K177A). pCF146 was cleaved with NdeI and re-ligated, generating pCF173, in which nucleotides 870 to 1050 were eliminated from the coding region of *safA.* Plasmid pCF138 was cleaved with HindIII and XhoI, releasing a 458 bp fragment that contains the segment of the *safA* region coding for the K318A substitution. This fragment was inserted into the corresponding sites of pCF173, to create pCF174 in which the C30 sequence was regenerated and now harbors the point mutations leading to the K177A and K318A substitutions. Plasmids pCF138, pCF146, or pCF174 were introduced into *E. coli* BL21(DE3) generating strains AH10529, AH10530 and AH10531, respectively.

**SafA^N21^**: The region from the 5’ end of the *safA* open reading frame until codon 164 was PCR-amplified with primers safA490Fw and safA980Rv from pCF70 (23) and the resulting fragment inserted between the XhoI and NcoI sites of pET16b to yield pBG2.

All constructs designed for the overproduction of SafA^FL^, SafA^N21^, C30 and its variants have a C-terminal His_6_-tag.

**Overproduction and purification of SafA^FL^, SafA^N21^, C30 and C30 variants.** *E. coli* BL21(DE3) was transformed with the following plasmids (protein to be overproduced indicated between parenthesis): pCF70 (SafA^FL^-His_6_), pCF68 (C30^WT^-His_6_), pCF174 (C30^K to A^-His_6_), pKA17 (C30^N ter^-His_6_), pKA18 (C30^C ter^-His_6)_, pKA32 (C30^C to S^-His_6_), pKA33 (C30^K to A/C to S^-His_6_), and pBG2 (SafA^N21^-His_6_) to create strains AH10326, AH10325, AH10531, AHEC1428, AHEC1429, AHEC1308, AHEC1309 and AHEC1101, respectively (see **S3_Table**). The strains were grown at 37°C, 150 rpm, for 18h, in an auto-induction medium (25) supplemented with 30 μg/ml kanamycin and with ampicillin 100 µg/ml in the case of AHEC1101.The cells were harvested by centrifugation (10 min at 3,500 x *g*, 4°C). Cells carrying C30 variants were resuspended at 1/10 of the culture volume in lysis buffer (50 mM NaH_2_PO_4_, 0.5 M NaCl, 10 mM Imidazole, pH 8.0). Cells carrying SafA^FL^ or SafA^N21^ were also resuspended in 1/10 of the culture volume of lysis buffer but supplemented with 2 mM DTT. Cells were lysed in a French Press cell (at 19000 lb/in^2^) and lysates were cleared by centrifugation (≥30 min at 27200 x *g*, 4°C). All the proteins were purified from the cleared lysates by Ni^2+^-NTA affinity chromatography; the columns were pre-equilibrated with 10x column volume lysis buffer, and washed with 10 ml lysis buffer (supplemented with 2 mM DTT in the case of SafA^FL^ and SafA^N21^) containing 10% glycerol. An imidazole step gradient (60, 150 and 250 mM in the case of SafA^FL^ and C30; 40, 100 and 300 mM in the case of SafA^N21^; each in 5 ml lysis buffer) was applied to elute each of the proteins of interest. SafA^FL^, C30 and its variants eluted at 250 mM imidazole, and the SafA^N21^ eluted at 100 mM imidazole. The purified fractions of C30 where dialyzed overnight at 1/1000 of the protein fraction volume against 0.1 M Tris-HCl, pH 8.0 (when samples were used for PAGE analysis, only) or against 0.1 M Tris-HCl, 0,15 M NaCl, pH 8.0 (when samples were used for SEC and SAXS studies) using a 10 kDa cutoff *SnakeSkin* membrane (from Pierce). The same protocol was used to dialyze SafA^FL^ and SafA^N21^, except that the dialysis buffer was supplemented with 2 mM DTT. The dialyzed fractions were split into aliquots, frozen in liquid nitrogen and stored at −80 °C until further use.

**Overproduction and purification of Tgl and Tgl^R185A^. Tgl^R185A^:** pLOM4 is a pET30a derivative carrying the WT *tgl* gene (56) (Fernandes et al., 2018). Quick-Change site-directed mutagenesis was used to substitute the Arg185 codon in pLOM4 with an Ala codon, using primer pair *tgl*-R185AD and *tgl*-R185AR and generating pKA22. Both pLOM4 and pKA22 allow for the production of proteins with a C-terminal His_6_-tag under the control of a T7*_lac_* promoter. The plasmids were introduced into BL21 (DE3) generating strains AH10178 and AHEC1451, respectively. The overproduction, purification, dialysis and storage steps are the same as described above for the SafA proteins except that the cells lysis was performed in buffer A (2mM CAPS pH 10.0, 0.5M NaCl, 10mM imidazole, pH 8.0) (C. G. Fernandes et al., 2018; Plácido et al., 2008).

***B. subtilis* expressing *safA* alleles at *amyE*.** The *safA* alleles of interest were introduced at the *amyE* locus of the previously described *ΔsafA* in-frame deletion mutant AH10297 (23,40), as follows: ***safA-C30^K to A^*:** plasmid pCF75, a pMLK83 derivative designed to insert *safA^WT^* at the *amyE* locus (C. G. Fernandes et al., 2018), was used as the template to introduce mutations coding for the K318A substitution or others (as specified below) via site directed mutagenesis using primers *safA*-K318A D and *safA*-K318A R; this generated pCF136. Plasmid pCF136 was subject to mutagenesis with primers *safA*-K177A D and *safA*-K177A R, generating pCF163 (which carries the *safA-C30^K177/K318 to A^* allele, referred to as *safA-C30^K to A^* for simplicity). This final plasmid was used to transform AH10297, complementing *ΔsafA* with *safA-C30^K to A^* at *amyE* and producing strain AH10493.

***safA-C30^C to S^*:** pCF73 (23) was used as template to introduce a point mutation causing the C65 to S substitution via site directed mutagenesis, using primers CtoS-65-Fw and CtoS-65-Rev. This generated pKA34, which carries the *safA-C30^C65 to S^* allele; PCR mutagenesis with primer pair CtoS-51-Fw and CtoS-51-Rev and pKA34 as the template, created pKA35 (carrying *safA-C30^51, C65 to S^*). Plasmid pKA35 was used as the template for a third round of site-directed mutagenesis, using primer pair CtoS_160/162_Fw and CtoS_160/162_Rev, to produce alleles coding for substitution of both residues, C160 and C162, with S residues. This produced pKA36, carrying *safA-C30^C51, C65, C160, C162 to S^* (C214, 228, 323, and 325 according to SafA^FL^ numbering); digestion of pKA36 with XhoI and BamHI released a 1416 bp fragment containing the promoter, the *safA* coding and terminator regions, which was inserted into pMLK83 to produce pKA46. Plasmid pKA46 was then used to transfer the *safA-C30^C51, C65, C160, C162 to S^* allele (*safA*-C30*^C to S^* for simplicity) to the *amyE* locus of AH10297, producing strain AH10623.

***safA-C30^K to A/C to S^*:** plasmid pCF163 (above), carrying *safA-C30^K to A^*, was used as the template for site directed mutagenesis using primer pair CtoS-51-Fw and CtoS-51-Rev, to produce the *safA-C30^K to A/C51 to S^*. This generated pKA47 which was in turn used as a template for a third round of site directed mutagenesis to give the *safA-C30 ^K to A/C51, C65to S^* allele, in pKA48. A third site directed mutagenesis PCR was performed using pKA48 as the template and primer pair CtoS_160/162_Fw and CtoS_160/162_Rev to produce, *C30 ^K to A/C51, C65, C160, C162 to S^*, in pKA49. This plasmid carries *safA-C30* with all K and C residues replaced by A or S residues, respectively (*safA-C30^K to A/C to S^*, for simplicity). Finally, pKA49 was used to transfer the *safA-C30^K to A/C to S^* allele to the *amyE* locus of the *ΔsafA* mutant AH10297, generating strain AH10624.

**Introduction of *safA-yfp* in a *tgl* deletion mutant:** To construct strains expressing *safA-yfp* in a *Δtgl* background, chromosomal DNA was extracted from strain AH10169 and used to transform AH10487 (23) with selection for Neo^R^ and Sp^R^, generating AH10797.

**Microscale Thermophoresis.** C30 or C30^C to S^ were used at various concentrations starting with 10 µM; Tgl was used at a concentration of 50 nM in 100 mM Tris-HCl, pH 8.0. To avoid aggregates, Tween was added to a final concentration of 0.05%; the samples (both the labelled protein and the ligands) were centrifuged at 14.000 rpm for 10 min at room temperature and the supernatant used in the assays. Assays were carried out on a Monolith instrument and data were analyzed with the MO.Affinity Analysis software v2.3both from NanoTemper (Munich, Germany).

**Structural Bioinformatics.** The structure of SafA^FL^ was predicted using AlphaFold3 (https://alphafoldserver.com/) based on its protein sequence. The resulting model was visualized and analyzed using ChimeraX (https://www.cgl.ucsf.edu/chimerax/). To identify disordered regions within the protein, we applied the deep-learning-based predictor Metapredict. We plotted the disordered propensity against the per-residue predicted local distance difference test (pLDDT) score. Additionally, we calculated key biophysical parameters such as hydrophobicity, net charge per residue (NCPR), fraction of charged residues (FCR), the partition coefficient of charges (κ), and diagram-of-states classification using the CIDER tool (Holehouse et al., 2017). To infer potential intermolecular interactions driven by the intrinsically disordered C30 region, we used FINCHES (31,32).

## SUPPORTING RESULTS AND DISCUSSION

### C30 is a proxy for the cross-linking of SafA^FL^ by Tgl

We found in this work that C30 forms large complexes that are cross-linked by Tgl. In earlier work, we showed that the activity of Tgl decreased with the concentration of substrate because Tgl became part of the cross-linked products (25). SafA^FL^ also forms large oligomeric complexes. To examine how Tgl cross-linked SafA^FL^, various concentrations of the purified protein (1.5, 3, 6 and 12 µM) were incubated with a fixed concentration of Tgl (4 μM, **S3A_Fig**). Samples were taken over time, for up to 120 min, and resolved by SDS-PAGE under reducing conditions to monitor the progression of the cross-linking reaction. The disappearance of the monomeric form of SafA^FL^ (which runs at about 50 kDa; red arrowhead in **S3A_Fig**) represents the cross-linking activity of Tgl. High molecular weight products of SafA^FL^, generically designated (SafA^FL^)_n_, that run above the 250 kDa marker band, were only detected at concentrations of the substrate higher than 3 µM (**S3A_Fig**). However, even at a concentration of 1.5 µM, the disappearance of SafA^FL^ could be easily quantified (**S3A_Fig**). The highest activity of Tgl was found for the lowest concentration of SafA^FL^ (1.5 µM) and the activity decreased with the increasing concentration of the substrate, with the highest drop from 6 to 12 µM (**S3B_Fig**). These results are consistent with the previous observations on the cross-linking of C30 by Tgl and suggest that Tgl becomes part of the cross-linked products jamming its ability to catalyze cross-linking reactions (**S3C_Fig** and **S4C_Fig**) (25). Consistent with this idea, the level of Tgl decreased significantly over time for reaction with 12 µM of SafA^FL^ (**S3C_Fig**). We reasoned that increasing the concentration of Tgl could overcome this “inhibitory” effect. To test this, at the “inhibitory” concentration of 12 µM, SafA^FL^ was incubated with increasing concentrations of Tgl (**S4A_Fig**). While very low activity was detected for concentrations of Tgl equal to 0.5, 1 and 2 µM the activity increased significantly at a concentration of 4 µM (**S4A_Fig** and **S4B_Fig**). Moreover, at 4 µM the level of Tgl decreased much less over time when compared to the lower concentrations of enzyme tested (**S4C_Fig**). In all, these results show that C30 and SafA^FL^ behave similarly (25).

### C30 is the only region of SafA^FL^ cross-linked by Tgl

Translation of the *safA* mRNA leads to the formation of 3 proteins, SafA^FL^, C30 and N21 (15). SafA^N21^ is a 21 kDa protein that comprises the N-terminal LysM domain of SafA^FL^, region A and the linker, up to Met codons 161 or 164 (15) (**Fig 1A** and **1B**). We wanted to determine whether Tgl cross-links all the domains or regions of SafA^FL^ or if cross-linking is restricted to the C30 domain. With that purpose, we examined the cross-linking of purified SafA^N21^ by Tgl. As for SafA^FL^, increasing concentrations of SafA^N21^ were incubated with Tgl for up to 120 min, and samples were collected throughout the experiment and resolved by SDS-PAGE in reducing conditions to monitor the formation of cross-linked products (**S5A_Fig**). For the highest concentrations of SafA^N21^ used, 6 and 12 µM, a very faint band accumulated that could correspond to a dimer (SafA^N21^_2_) and no high molecular weight forms were detected (**S5A_Fig**). We have not quantified the disappearance of the monomeric form of SafA^N21^. We conclude that at least *in vitro*, cross-linking of SafA^N21^ by Tgl does not occur or is very inefficient. It follows that *in vitro* SafA^FL^ is cross-linked mainly if not exclusively through its C30 domain (**5B_Fig**).

### Mapping the disulfide bonds in C30

To identify the disulfide bonds in C30, we performed a LC-MS peptide mapping analysis of the non-reduced protein after the digestion with chymotrypsin. The identification of disulfide bond Cys214-Cys228 and of Cys323-Cys325 is shown in **Fig 4D** and **S6_Fig**, respectively. For Cys214-Cys228, we observed MSMS evidence for the presence of the disulfide bond in the peptide

HYPAHFVP**C^214^**PVPVSPILPGSGL**C^228^**YPY. The MSMS ions highlighted in **Fig 4D** correspond to the internal fragments with the disulfide bond (HYPAHFVP**C^214^[*]**PVP/GSGL**C^228^[*]**YPY; HYPAHFVP **C^214^[*]**PVPVSP/GSGL**C^228^[*]**YPY; HYPAHFVP**C^214^[*]**PVPVSPI/PGSGL**C^228^[*]**YPY). In agreement with the presence of the disulfide bond, we only detected b_1-7_ ions and y_1-3_ ions. Similar results were observed for the chimiotryptic peptide VP**C^214^**PVPVSPILPGSGL**C^228^**YPY. The disulfide bond Cys323-Cys325 was detected in peptide ADHDD**C^323^**G**C^325^**DGDHQPY (represented in **Fig 4D**; spectrum in **S6_Fig**). The MSMS ions highlighted in **S6_Fig** correspond to the internal fragments with the disulfide bond (**S2_Table**). Similar results were obtained for the chimiotryptic peptide MPYADHDD**C^323^**G**C^325^**DGDHQPY.

### Tgl cross-linking of C30 in the pre-formed complexes

We sought to gain insight into how Tgl cross-links C30 in the pre-formed complexes. We compared the cross-linking of C30^WT^ by either Tgl^WT^ or by a form of the enzyme, Tgl^R185A^, which we have shown before to cross-link substrate proteins at a slower rate (25). Purified C30^WT^ was incubated with Tgl^WT^ or Tgl^R185A^, and samples were collected over time and analyzed by SDS-PAGE under reducing conditions. With Tgl^WT^, C30 was detected with the apparent sizes of a dimer, a trimer, and a tetramer from min 5 onwards, and these were progressively replaced until, from 30 min onwards, (C30_6_)_n_ is the predominant species detected (**S8A_Fig**). When C30^WT^ was cross-linked by Tgl^R185A^, all these species were detected but persisted for longer times and (C30_6_)_n_ only became the most prevalent species at 120 min, consistent with the slower kinetics of Tgl^R185A^ (**S8B_Fig**) (25). The [(C30_6_)_n_]_n_ species is only detected in the presence of Tgl^WT^ (**Fig 6A** and **S8A_Fig**); its formation may be slow explaining why it was not detected in the reaction with Tgl^R185A^ (**S8B_Fig**).

A dimer is the first species to accumulate in the presence of Tgl or Tgl^R185A^ (**S8_Fig).** We suggested above that a dimer is the basic, repeating unit in the self-assembly of the large C30 oligomers (see above; **Fig 3B**). In one model, all the three dimers in C30_6_, taken as an example of an oligomeric complex, would be intra-molecularly cross-linked, via K177, K318 or both, and only then intra-molecular cross-linking (between dimers) would ensue. This model predicts that only multiples of the mass of the dimer would be detected by SDS-PAGE, *i.e*., a trimer with a mass of approximately 90 kDa should not be detected. However, a species that migrates between the 75 kDa and 100 kDa markers is seen in the reaction with Tgl^R185A^ (**S8B_Fig**, overexposed 120 min lane). At the same time, we proposed that K177 may only form intermolecular cross-links while K318 may only form intramolecular bonds (see main text; see also **Fig 7F**). Thus, one model is that the cascade may start with the cross-linking of a dimer via K177, followed by cross-linking of this dimer to one monomer of another dimer through K318, then cross-linking of that dimer via K177, with the sequence repeated until all the C30 molecules in the oligomers are cross-linked (**S8D_Fig**).

## SUPPORTING INFORMATION

**S1_Fig.**
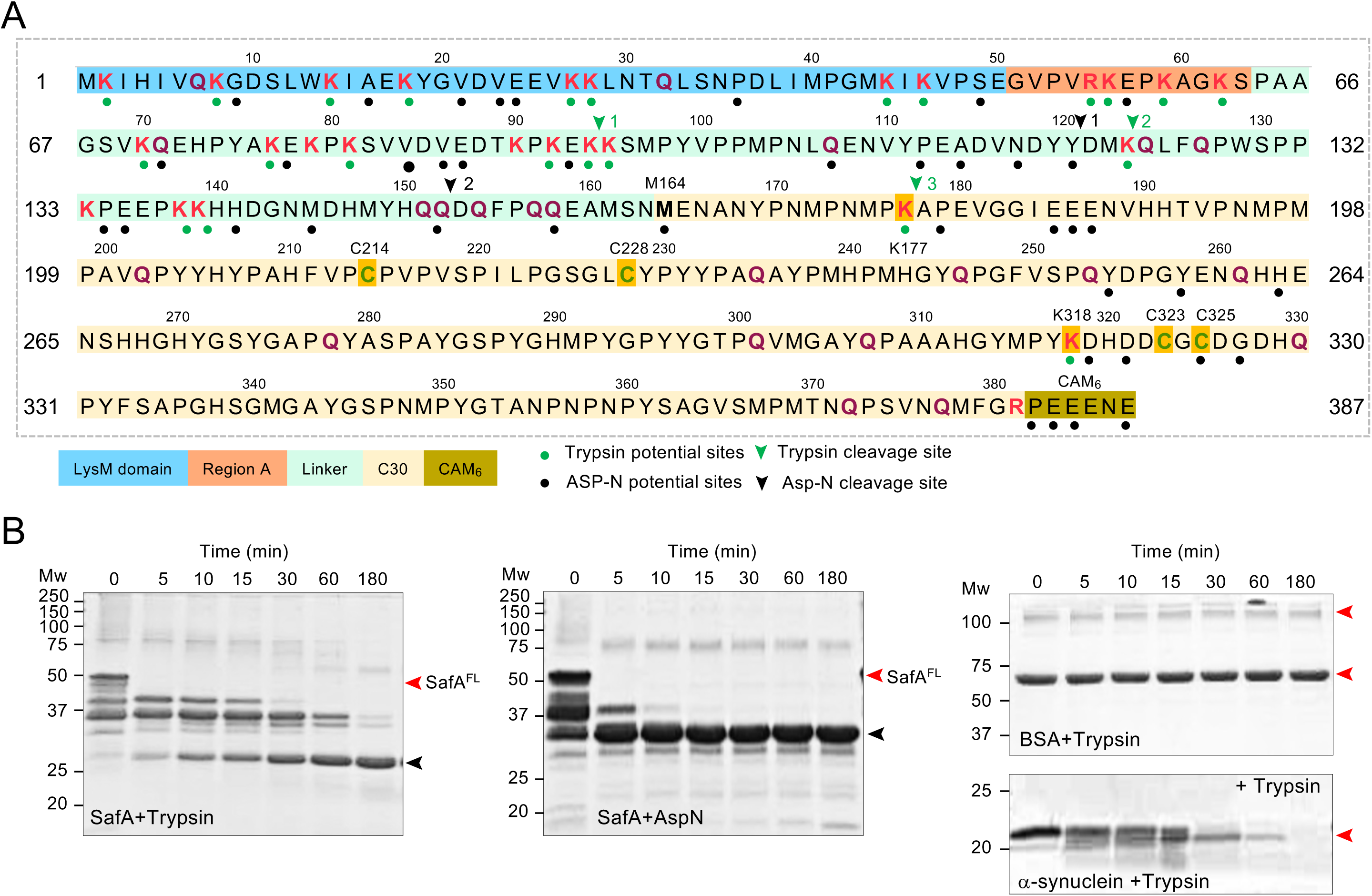
Primary structure and organization of SafA. **A**: The domains of SafA^FL^ are indicated: LysM, light blue; region A, light orange; the linker, light mint green; and C30, pale yellow. The 6 amino acid-long C-terminal acidic peptide (CAM_6_: PEEENE) is highlighted in mustard yellow. Relevant amino acids are highlighted as follows: Cys residues, green; Lys residues, reddish-pink; Gln residues, magenta. Potential cleavage sites for Trypsin and Asp-N are indicated by green or black dots, respectively and verified cleavage sites are shown by arrowheads with numbers indicating the order of cleavage. **B.** C30 is resistant to proteolysis. SafA^FL^ was incubated with either Trypsin or AspN (left and middle panels). Control experiments for trypsin digestion were performed using BSA or ⍺-sinuclein as substrates (right panels). The reactions were conducted for 180 min, samples were collected at the indicated time points and resolved by SDS-PAGE (15%) under reducing conditions and the gels stained with Coomassie solution. Gels run in parallel were transferred to PVDF membranes for N-terminal sequencing analysis. Red arrowheads mark the position of SafA^FL^ in the left and the middle panels and the forms of BSA and ⍺-sinuclein in the right panels. The black arrowhead marks the position of a stable proteolytic fragment of SafA^FL^ that contains part of C30. Mw markers (KDa) are shown on the left side of the panels.

**S2_Fig.**
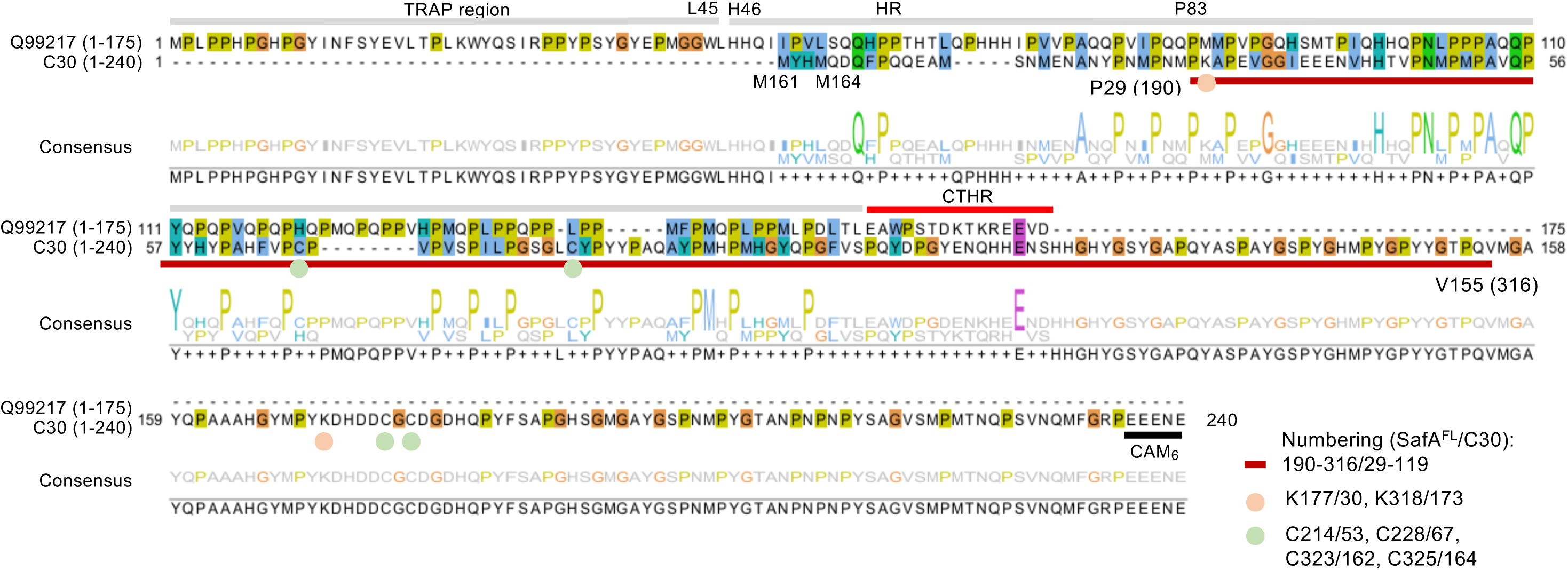
Similarity between C30 and human amelogenin. The figure shows the alignment of the amino acid sequences of human amelogenin (UniProt code: Q99217) and C30. The alignment was generated using Clustal W (www.ebi.ac.uk) and edited using JalView (https://www.jalview.org). The two possible starts for C30, M161 or M164, are indicated. The Lys (K177 and K318) and Cys (C214, C228, C323 and C325) residues in C30 are indicated, as well as the CAM_6_ acidic motif at its C-terminus (see also **Fig 2A**). The domain organization of amelogenin is indicated (lines above the sequence): the N-terminal hydrophilic, tyrosine-rich region (TRAP), the central hydrophobic region, rich in histidine, glutamine and proline residues (HR) (grey lines above the sequence) and the C-terminal hydrophilic region (CTHR). The brown line below the C30 sequence represents the stretch of residues thought to include self-interacting motifs (positions 191-316 of SafA^FL^ or P29 to V155 of C30) as defined in **Fig 2B** and 2**E** (see the main text for details).

**S3_Fig.**
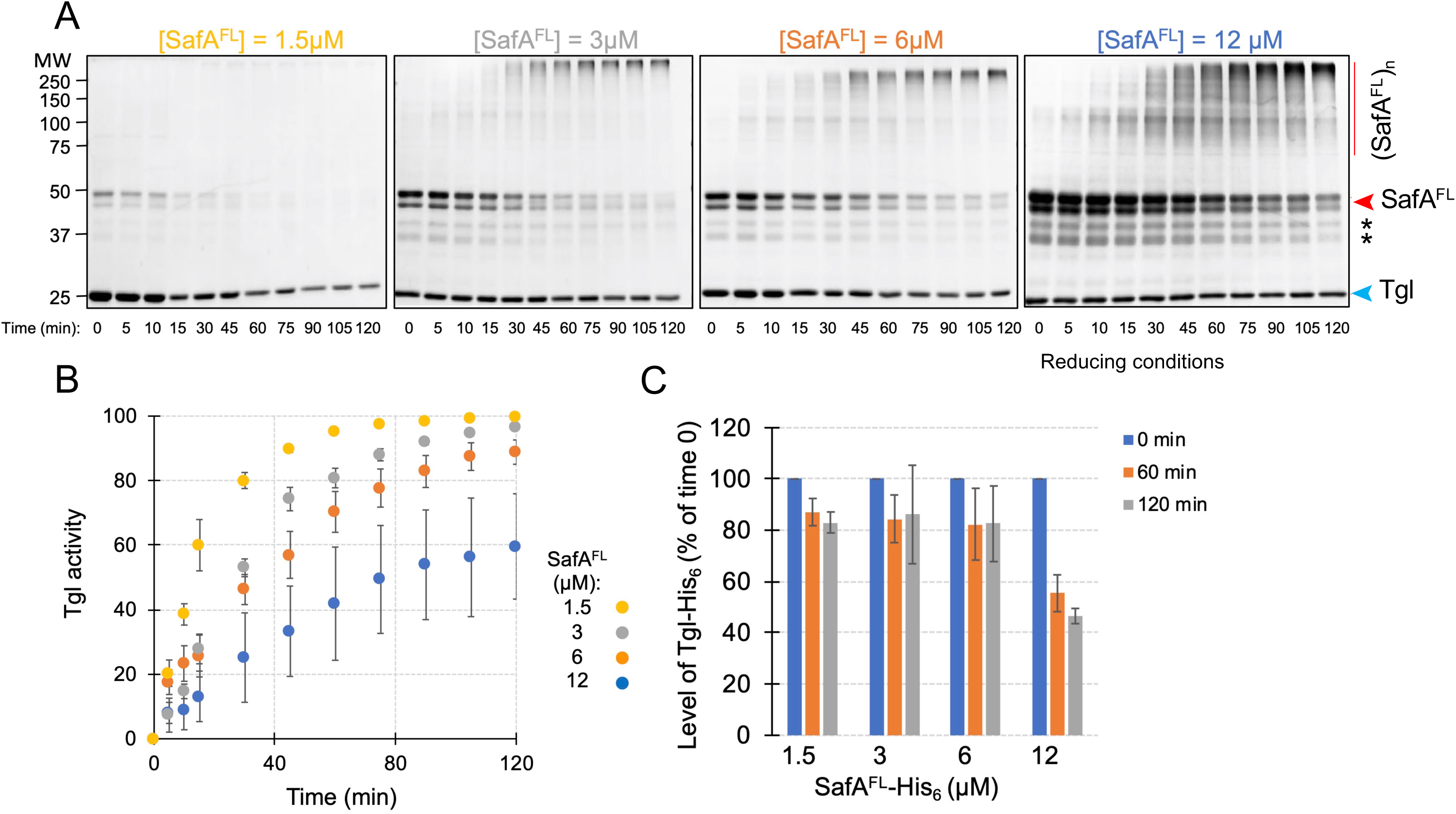
Tgl activity decreases with the concentration of SafA^FL^. **A**: The indicated concentrations of SafA^FL^ were incubated with Tgl (4 μM) at 37°C; samples were collected at the indicated time points and resolved by SDS-PAGE (10%) under reducing conditions. Red and blue arrowheads mark the position of SafA^FL^ and Tgl, respectively. The asterisk denotes likely degradation products of SafA^FL^. The red line shows the position of cross-linked products of SafA^FL^, [(SafA^FL^)n]. Mw markers (kDa) are shown on the left side of the panels. Gels were stained with Coomassie. **B**: Tgl activity was estimated by measuring the loss of SafA^FL^ over time using Image J. **C**: Decrease in the level of Tgl-His_6_, at a fixed concentration of 4 μM, at the indicated times after the onset of the reaction (time 0), as a function of the concentration of SafA^FL^.

**S4_Fig.**
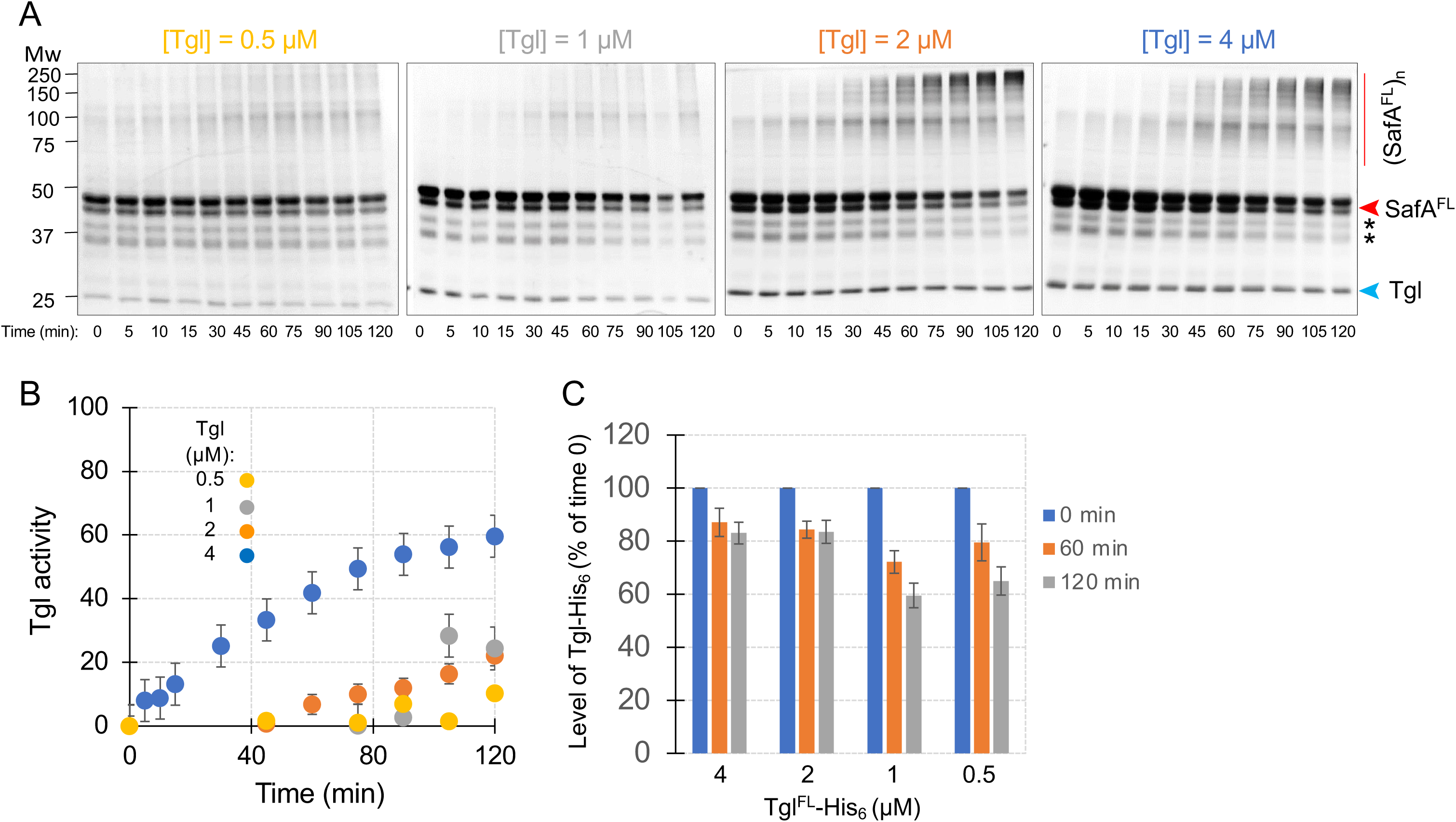
Cross-linking of SafA^FL^ increases with the concentration of Tgl. **A**: SafA^FL^ (12 μM) was incubated with different concentrations of Tgl, as indicated, at 37°C. Samples were collected at the indicated time points and resolved by SDS-PAGE (10%) under reducing conditions and the gels were stained with Coomassie. Red and light blue arrowheads mark the position of SafA^FL^ and Tgl, respectively. The asterisk denotes likely degradation products of SafA^FL^. The red line shows the position of cross-linked products of SafA^FL^, denoted as [(SafA^FL^)_n_]. Mw markers (in kDa) are shown on the left side of the panels. **B**: Tgl activity was measured and normalized as a function of the loss of SafA^FL^ over time, using Image J. **C**: Decrease in the level of Tgl at the indicated time points, as a function of the concentration of Tgl, for a fixed concentration of SafA^FL^ (12 μM).

**S5_Fig.**
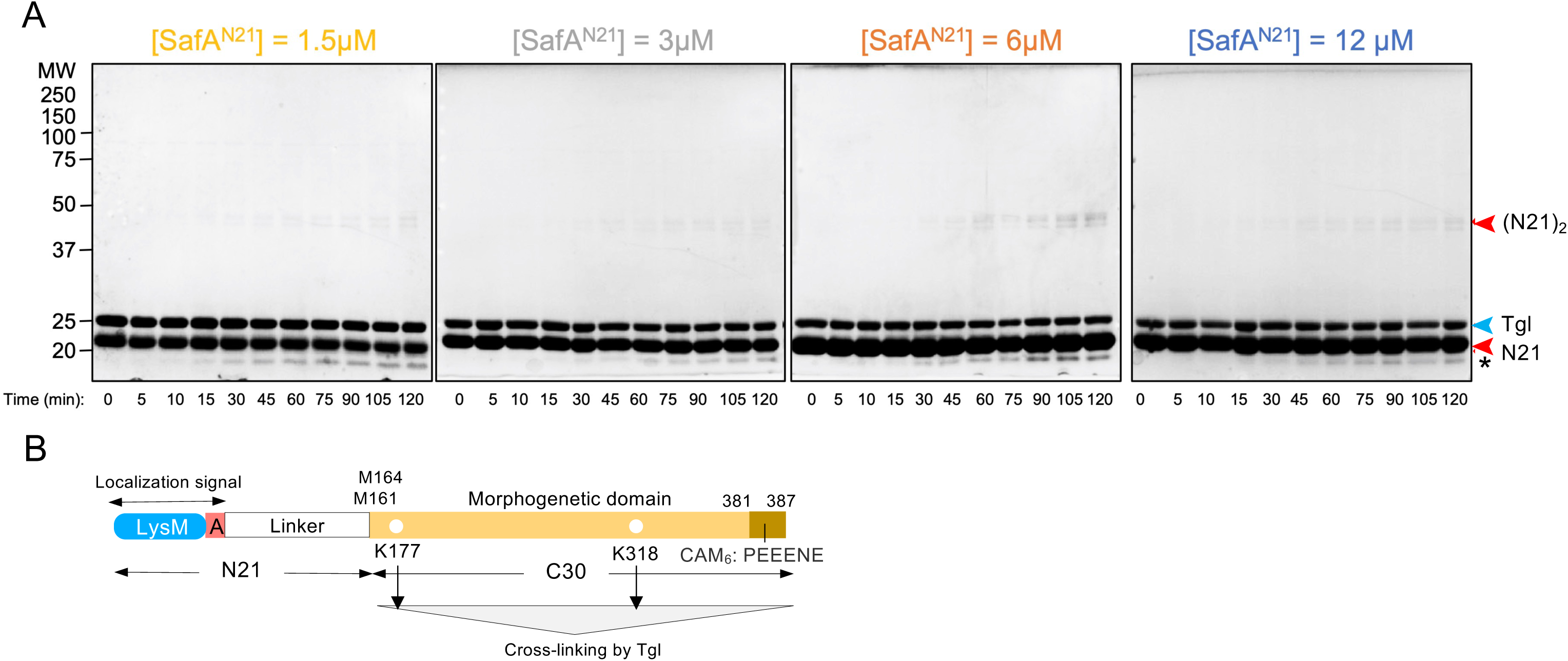
SafA^N21^ is not efficiently cross-linked by Tgl. **A**: SafA^N21^ was incubated, at four different concentrations with a fixed concentration of Tgl (4 μM) at 37°C. Samples were collected at the indicated time points, resolved by SDS-PAGE (10%) under reducing conditions and the gels stained with Coomassie. Red and light blue arrowheads mark the position of SafA^N21^ and Tgl, respectively. The asterisk mark the position of a possible intramolecular cross-linked species. Mw markers (kDa) are shown on the left side of the panels. **B**: Domain organization of SafA^FL^. At least *in vitro*, cross-linking of the protein by Tgl seems restricted to the C30 domain.

**S6_Fig.**
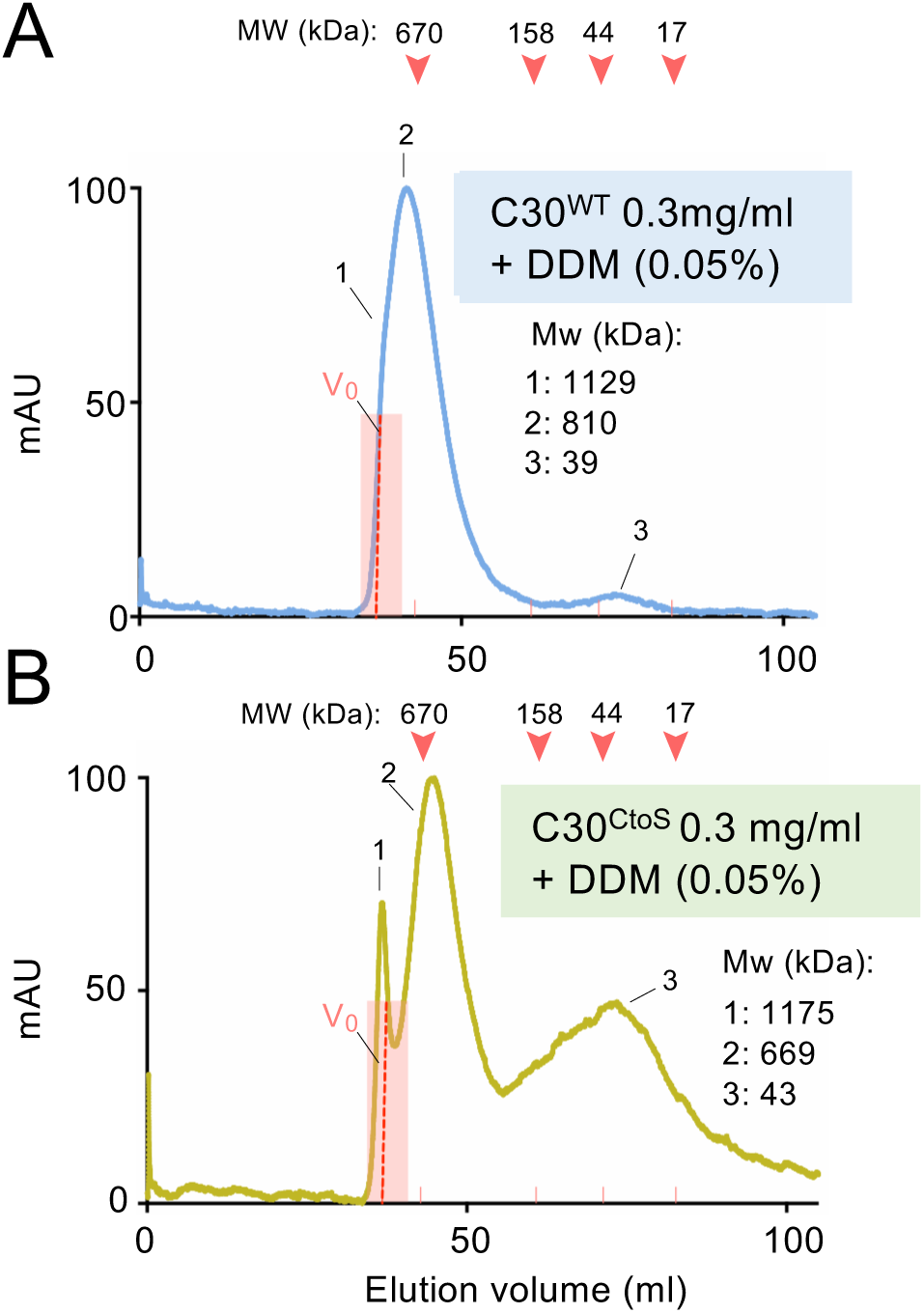
Effect of DDM on the complexes formed by C30. C30^WT^ (**A** and **B**) and C30^C to S^ (**C** and **D**) were loaded at a concentration of 0.3 mg/ml on a Sephacryl S-300 SEC column in the absence (**A** and **C**) or in the presence (**B** and **D**) of 0.05% DDM. The protein and DDM concentrations mimic those used for the Blue Native PAGE analysis. The position of the peaks is identified for each chromatogram and an estimation of the MW is shown. The position of MW markers is shown by the light red arrowheads.

**S7_Fig.**
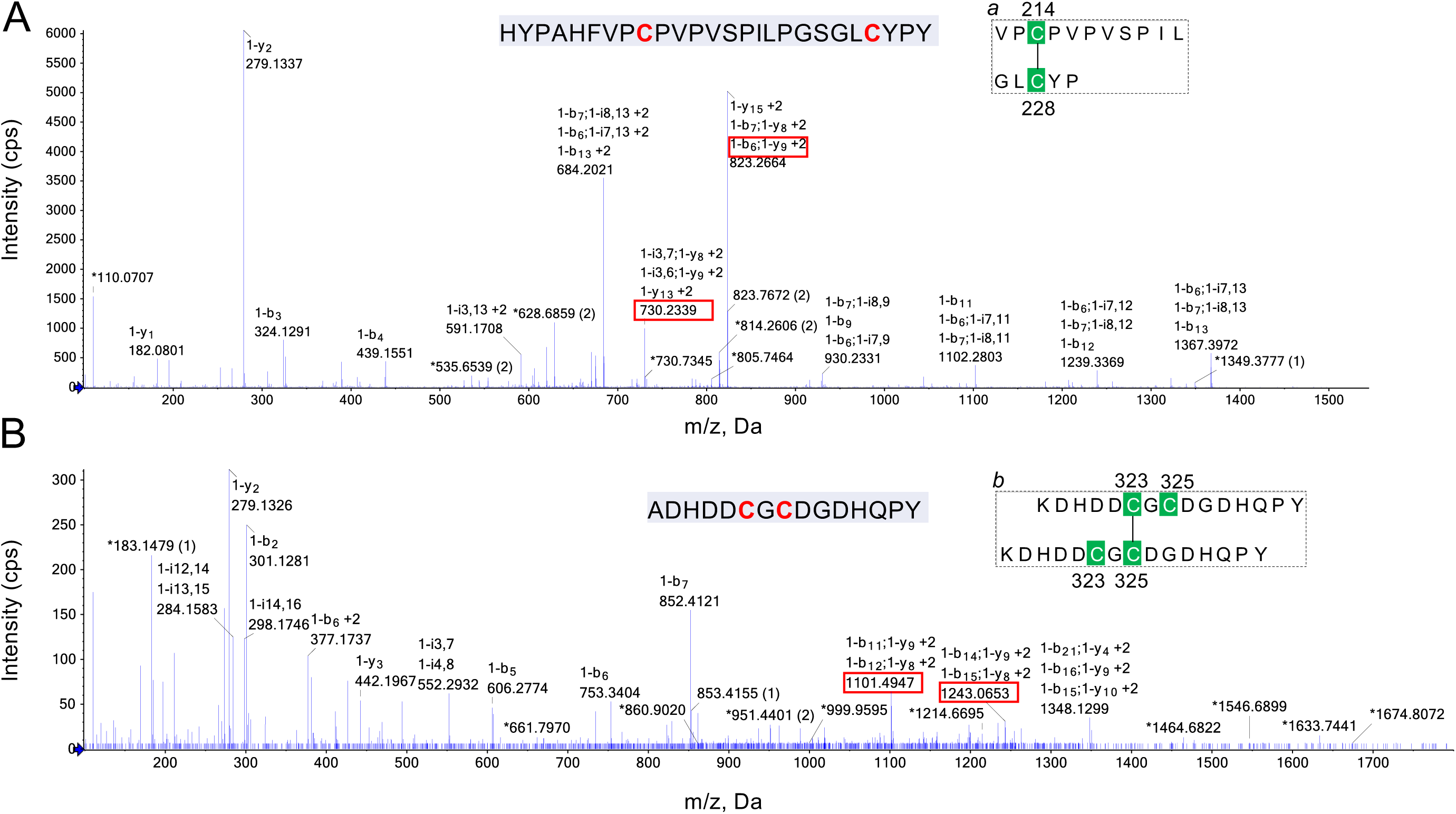
Identification of the Cys214-Cys228 and Cys323-Cys325 cross-links. **A** and **B**: LC-MSMS analysis of disulfide bond Cys214-Cys228 and Cys323-Cys325 (numbering is from the N-terminal of SafA^FL^). The MSMS fragmentation pattern is show. The highlighted ions identify the Cys214-Cys228 and Cys323-Cys325 disulfide bonds. See also **Fig 4D**.

**S8_Fig.**
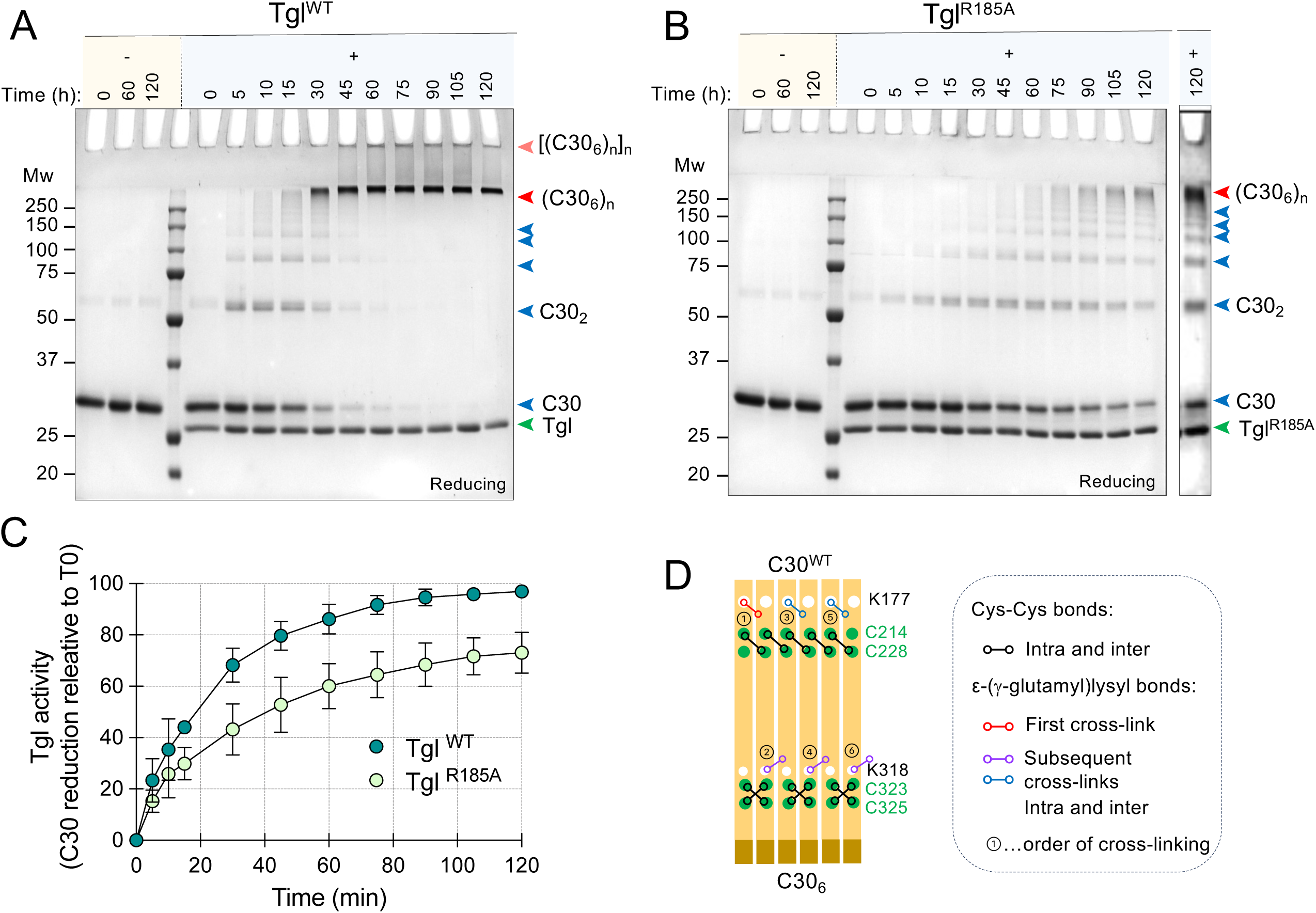
Cross-linking within the C30 oligomers. Purified C30 was incubated for up to 120 min in the absence (“-“) or in the presence Tgl^WT^ or its variant Tgl^R185A^ (“+”). Samples withdrawn at the indicated time points (in min) were resolved by reducing SDS-PAGE (10%). In **A** and **B**, the blue arrowheads show the position of C30 and multimeric forms of the protein; the red arrowheads show the positions of (C30_6_)_n_ and [(C30_6_)_n_]_n_. The green arrowhead shows the position of Tgl. Gels were stained with Coomassie. **C**: Comparison of the activity of Tgl^WT^ and Tgl^R185A^, measured by the rate of disappearance of C30. **D**: a model for the cross-linking of C30 units within the oligomer, taken as an example. The model considers the possibility that K177 may only be involved in intermolecular cross-linking (between dimers) whereas K318 may only form intramolecular cross-links (within a dimer). The first cross-linking event involves K177 (1); then, a cross-link forms via K318 (2) followed by another cross-link involving K177 (3), until all the C30 units in the oligomer are cross-linked. The model accounts for the detection of a C30 trimer by SDS-PAGE.

**S9_Fig.**
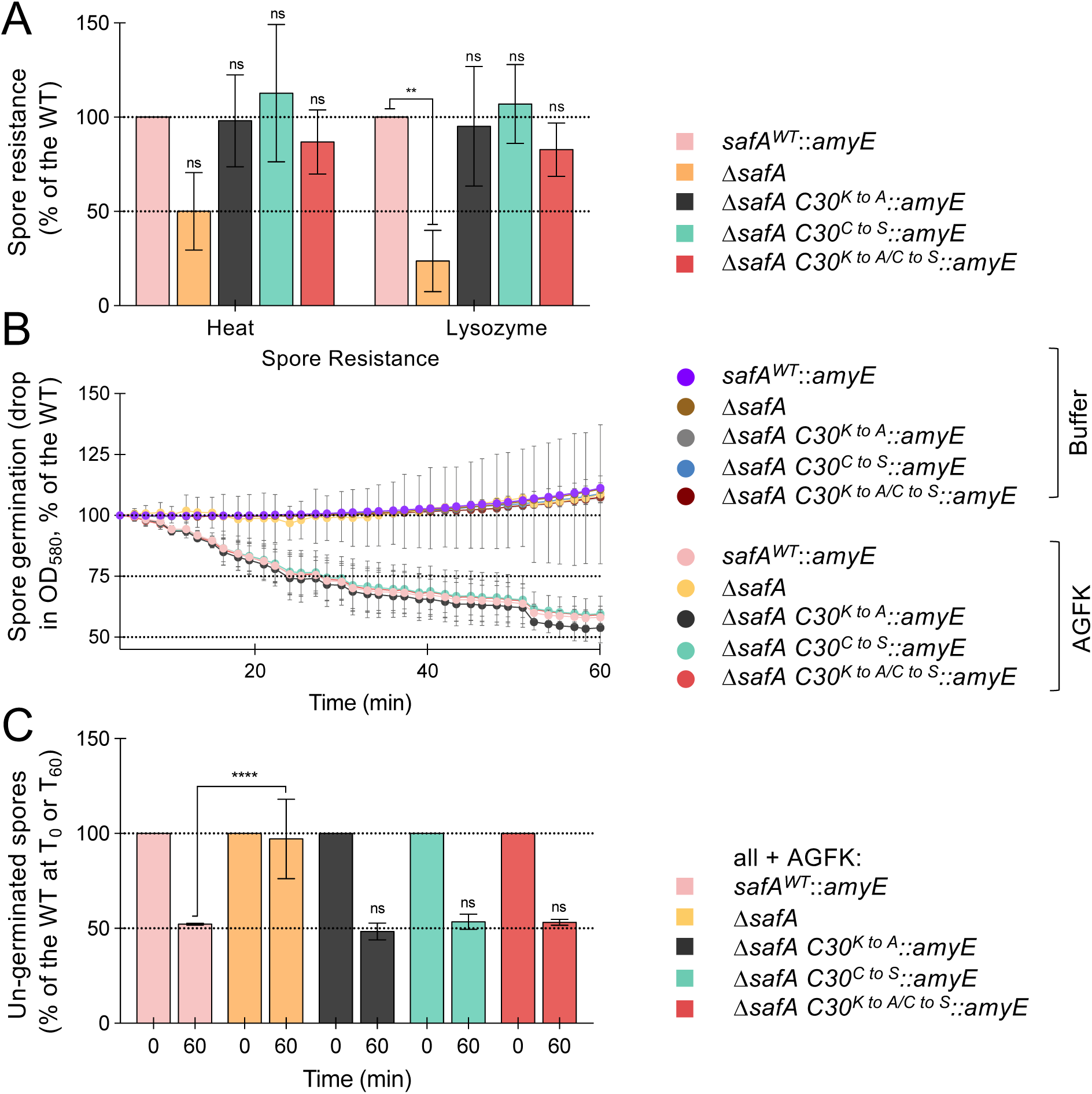
Spore properties. A: Heat and Lysozyme resistance of *safA* and *C30* spores. Spores were density gradient purified from 24h sporulating cultures of a Δ*safA* mutant or strains bearing either the WT *safA* allele or point mutations in *C30* inserted at the non-essential *amyE* locus in a *safA* in-frame deletion mutant. Aliquots of the spore suspensions were serially diluted and plated on LB agar plates before and after incubation with lysozyme or at 80°C (for 20 min). After overnight incubation, the survivors were enumerated. The data shown are the mean and standard deviation (SD) of the results from three independent experiments and are normalized to the reference strain (M*safA* with *safA*^WT^ at *amyE*). **B: Kinetics and extent of spore germination:** spores of the indicated strains were density gradient purified and resuspended in a buffer. Germination was induced by adding a mixture of asparagine, glucose, fructose and KCl (AGFK) and followed by monitoring the drop in the OD_580_ for up to 1 hour. Control cultures were incubated in buffer, with no addition of AGFK, for the same duration. **C**: shows the extent of germination for the indicated strains, measured before adding the germinant mixture and 60 min thereafter, normalized to the reference strain. In **A, B** and **C**: the data shown are the mean and standard deviation (SD) of the results from three independent experiments. The data in **A** and **C** are normalized to the reference strain. Statistical analysis used a one-way ANOVA followed by Dunnett’s multiple comparison test (⍺ = 0.05). Statistical significance(*P* < 0.05) is indicated by asterisks **, P < 0.05; ****, P < 0.0001, and “ns” for non-significant differences.

## SUPPORTING TABLES

**S1_Table.**
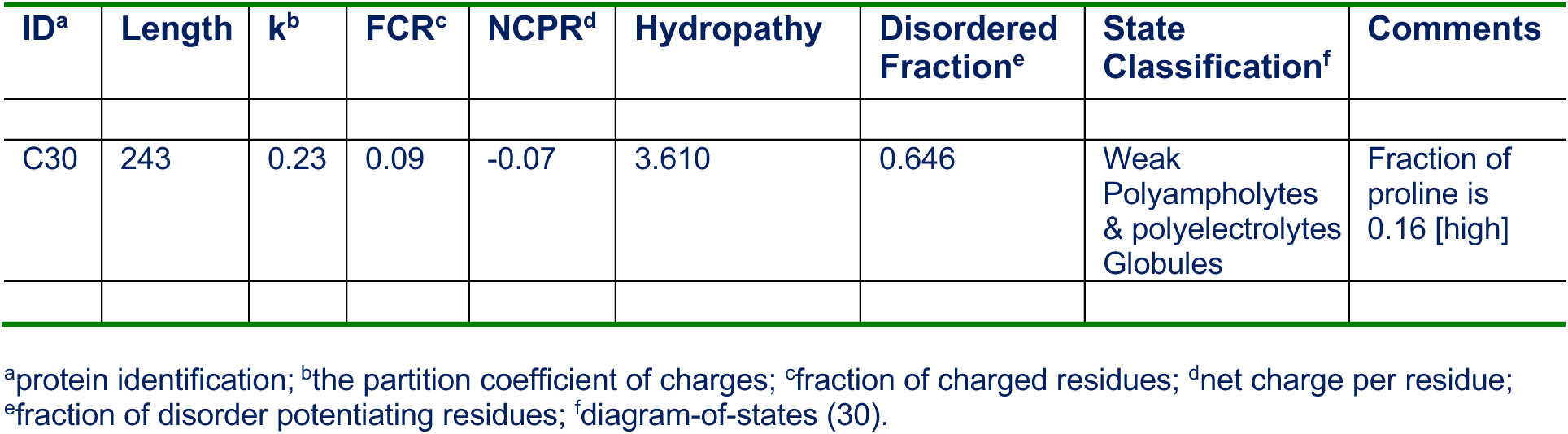
SafA properties.

**S2_Table.** MSMS data of peptide ADHDDC^323^GC^325^DGDHQPY carrying disulfide bond Cys323-Cys325.

| Fragment | Type | Observed m/z |
| --- | --- | --- |
| HDDC[*1]GC[*1]DGDHQPYP | y | 730.23395 |
| HDDC[*1] / GC[*1]DGDHQPYP | b,2y |  |
| HDDC[*1]G / C[*1]DGDHQPYP | b,2y |  |
| ADHDDC[*1] / GC[*1]DGDHQPYP | i | 823.26638 |
| ADHDDC[*1]G / C[*1]DGDHQPYP | i |  |
| ADHDDC[*1] / GC[*1]D | 2b,y | 930.23305 |
| ADHDDC[*1]GC[*1]D | b |  |
| ADHDDC[*1]G / C[*1]D | 2b,y |  |
| ADHDDC[*1]G / C[*1]DGD | 2b,y | 1102.28032 |
| ADHDDC[*1] / GC[*1]DGD | 2b,y |  |
| ADHDDC[*1]GC[*1]DGD | b |  |
| ADHDDC[*1]GC[*1]DGDH | b | 1239.33685 |
| ADHDDC[*1]G / C[*1]DGDH | 2b,y |  |
| ADHDDC[*1] / GC[*1]DGDH | 2b,y |  |
| ADHDDC[*1]GC[*1]DGDHQ | b | 1367.3972 |
| ADHDDC[*1]G / C[*1]DGDHQ | 2b,y |  |
| ADHDDC[*1] / GC[*1]DGDHQ | 2b,y |  |

**S3_Table.** Bacterial strains used in this study.

| Strain | Relevant Properties* | Origin/Reference |
| --- | --- | --- |
| *E. coli* |  |  |
| DH5α | F– Φ80*lacZΔM15 Δ(lacZYA-argF) U169 recA1 endA1 hsdR17* (rK–, mK+) *phoA supE44 λ– thi-1 gyrA96 relA1* | Invitrogen |
| BL21 (DE3) | *fhuA2 [lon] ompT gal (λ DE3) [dcm] ΔhsdS λ DE3 = λ sBamHlo ΔEcoRI-B int::(lacI::PlacUV5::T7 gene1) i21 Δnin5* | Novagen |
| AH10178 | BL21 (DE3) carrying pLOM4 | (56) |
| AH10325 | BL21 (DE3) carrying pCF68 | (59) |
| AH10326 | BL21 (DE3) carrying pCF70 | " |
| AH10531 | BL21 (DE3) carrying pCF174 | " |
| AHEC1101 | BL21 (DE3) carrying pBG2 | This study |
| AHEC1308 | BL21 (DE3) carrying pKA32 | " |
| AHEC1309 | BL21 (DE3) carrying pKA33 | " |
| AHEC1428 | BL21 (DE3) carrying pKA17 | " |
| AHEC1429 | BL21 (DE3) carrying pKA18 | " |
| AHEC1451 | BL21 (DE3) carrying pKA22 | " |
| *B. subtilis* |  |  |
| MB24 | *trpC2 metC3* (WT, Spo<sup>+</sup>) | (59) |
| AH10169 | *trpC2 metC3 tgl::sp* | (23) |
| AH10297 | *trpC2 metC3 ΔsafA ΔamyE::cm* | " |
| AH10487 | *trpC2 metC3 ΔsafA ΔamyE::safA-yfp* | (13) |
| AH10493 | *trpC2 metC3 ΔsafA amyE::safA<sup>K to A</sup>* | This study |
| AH10623 | *trpC2 metC3 ΔsafA ΔamyE::safA<sup>C to S</sup>* | " |
| AH10624 | *trpC2 metC3 ΔsafA ΔamyE::safA<sup>K to A, C to S</sup>* | " |
| AH10797 | *trpC2 metC3 tgl::sp ΔsafA ΔamyE::safA-yfp* | " |

\*Km<sup>R</sup>, Neo<sup>r</sup>, Cm<sup>r</sup>, Sp<sup>r</sup>: kanamycin, neomycin, cloranfenicol or spectinomycin resistance.

**S4_Table.**
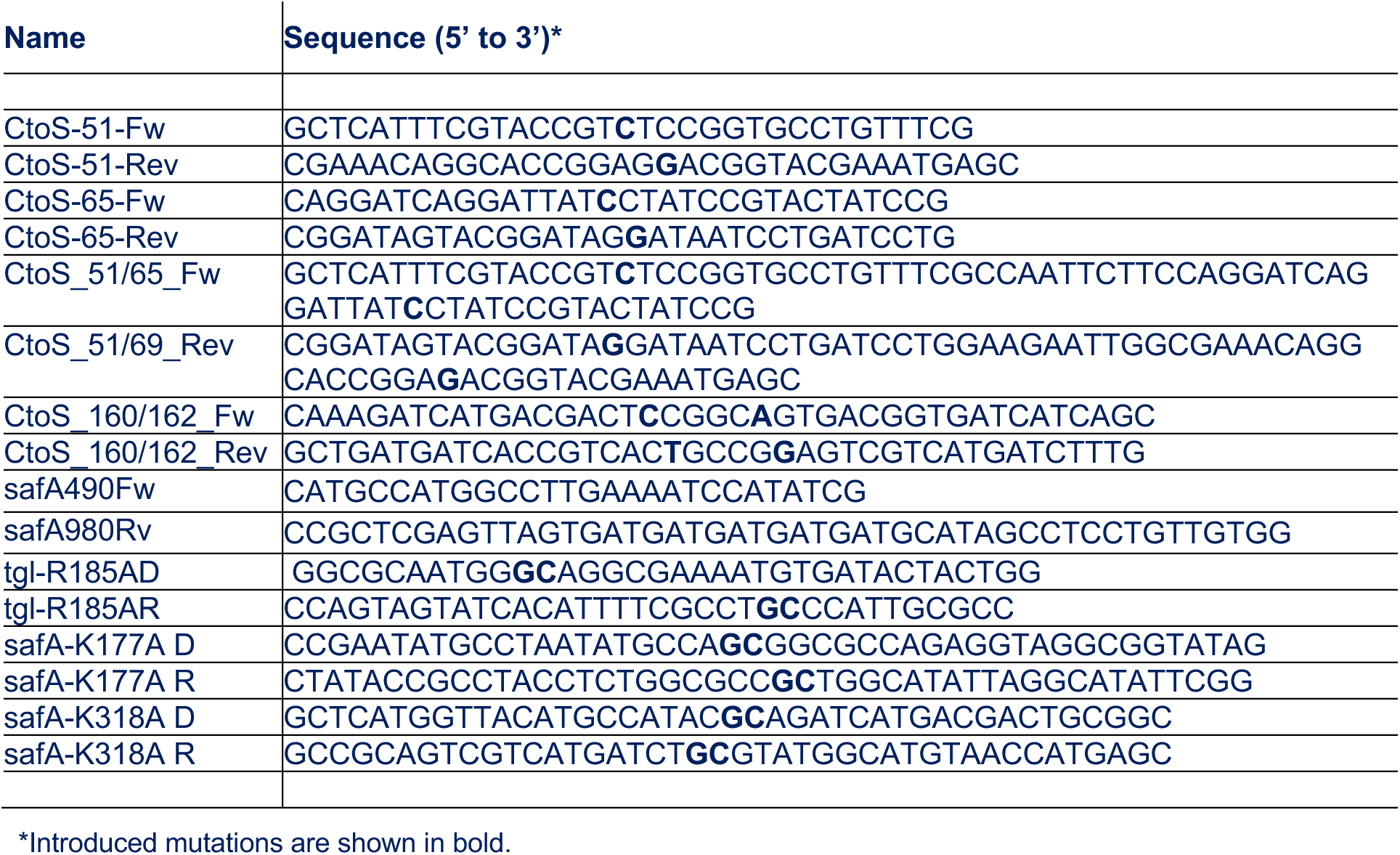
Oligonucleotide primers.

**S5_Table.** Plasmids used in this study.

| Plasmid | Relevant features | Origin/Reference |
| --- | --- | --- |
| pET16b | His<sub>6</sub>-tag fusion protein expression vector (Amp<sup>R</sup>)* | Novagen |
| pET28a+ | His<sub>6</sub>-tag fusion protein expression vector (Km<sup>R</sup>)* | " |
| pET30a+ | His<sub>6</sub>-tag fusion protein expression vector (Km<sup>R</sup>) | " |
| pMLK83 | Plasmid for marker replacement integration at *amyE* (Neo<sup>R</sup>)* | Lab collection |
| pLitmus29 | High-copy number vector *E. coli* (Amp<sup>R</sup>)* | NEB |
| pCF68 | C30<sup>WT</sup>-His<sub>6</sub> in pET28a+ (Km<sup>R</sup>) | (25) |
| pBG2 | SafA<sup>N21</sup>-His<sub>6</sub> in pET16b (Amp<sup>R</sup>) | " |
| pCF70 | SafA<sup>FL</sup>-His<sub>6</sub> in pET28a+ (Km<sup>R</sup>) | (59) |
| pCF73 | pLitmus 29 derivative carrying *safA* (Amp<sup>R</sup>) | " |
| pCF75 | pMLK83 derivative for insertion of *safA* at the *amyE* locus (Neo<sup>R</sup>) | " |
| pCF136 | pMLK83 derivative for insertion of *safA*-C30<sup>K318 to A</sup> at *amyE* (Neo<sup>R</sup>) | " |
| pCF149 | pMLK83 derivative for insertion of *safA*-*yfp* at *amyE* (Neo<sup>R</sup>) | (13) |
| pCF163 | pMLK83 derivative for insertion of *safA*-C30<sup>K to A</sup> at *amyE* (Neo<sup>R</sup>) | This study |
| pCF174 | C30<sup>K to A</sup>-His<sub>6</sub> in pET28a+ (Km<sup>R</sup>) | " |
| pLOM4 | Tgl<sup>WT</sup>-His<sub>6</sub> in pET30a+ (Km<sup>R</sup>) | (56) |
| pCF68 | C30<sup>WT</sup>-His<sub>6</sub> in pET28a+ (Km<sup>R</sup>) | " |
| pKA17 | C30<sup>N-ter</sup>-His<sub>6</sub> in pET28a+ (Km<sup>R</sup>) | " |
| pKA18 | C30<sup>C-ter</sup>-His<sub>6</sub> in pET28a+ (Km<sup>R</sup>) | " |
| pKA22 | Tgl<sup>R185A</sup>-His<sub>6</sub> in pET30a+ (Km<sup>R</sup>) | " |
| pKA32 | C30<sup>C to S</sup>-His<sub>6</sub> in pET28+ (Km<sup>R</sup>) | " |
| pKA33 | C30<sup>C to S, K to A</sup>-His<sub>6</sub> in pET28a+ (Km<sup>R</sup>) | " |
| pKA34 | pCF73 derivative carrying *safA*-C30<sup>C65 to S</sup> (Amp<sup>R</sup>) | " |
| pKA35 | pKA34 derivative carrying *safA*-C30<sup>C51 to S, C65 to S</sup> (Amp<sup>R</sup>) | " |
| pKA36 | pKA35 derivative carrying *safA*-C30<sup>C to S</sup> (Amp<sup>R</sup>) | " |
| pKA46 | pMLK83 derivative for insertion of *safA*-C30<sup>C to S</sup> at *amyE* (Neo<sup>R</sup>) | " |
| pKA47 | pCF163 derivative carrying *safA*-C30<sup>K to A, C51 to S</sup> (Neo<sup>R</sup>) | " |
| pKA48 | pCF163 derivative carrying *safA*-C30<sup>K to A, C51 to S, C65 to S</sup> (C30<sup>K to A, C to S</sup>) (Neo<sup>R</sup>) | " |
| pKA49 | pMLK83 derivative for insertion of *safA*- C30<sup>K to A, C to S</sup> at *amyE* (Neo<sup>R</sup>) | " |

\*Km<sup>R</sup>, Amp<sup>R</sup>, Neo<sup>R</sup>: kanamycin, ampicillin and neomycin resistance.

